# Saturating the eQTL map in *Drosophila melanogaster*: genome-wide patterns of cis and trans regulation of transcriptional variation in outbred populations

**DOI:** 10.1101/2023.05.20.541576

**Authors:** Luisa F. Pallares, Diogo Melo, Scott Wolf, Evan M. Cofer, Varada Abhyankar, Julie Peng, Julien F. Ayroles

**Affiliations:** Lewis-Sigler Institute for Integrative Genomics, Princeton University, Princeton, NJ, USA; Ecology and Evolutionary Biology Department, Princeton University, Princeton, NJ, USA; Friedrich Miescher Laboratory of the Max Planck Society, 72076 Tübingen, Germany

**Keywords:** Expression Quantitative Trait Loci, eQTL, *Drosophila melanogaster*, *gene reg*ulation, transcription

## Abstract

Decades of genome-wide mapping have shown that most genetic polymorphisms associated with complex traits are found in non-coding regions of the genome. Characterizing the effect of such genetic variation presents a formidable challenge, and eQTL mapping has been a key approach to understand the non-coding genome. However, comprehensive eQTL maps are available only for a few species like yeast and humans. With the aim of understanding the genetic landscape that regulates transcriptional variation in *Drosophila melanogaster, we devel*oped an outbred mapping panel in this species, the *Drosophila* Outbred Synthetic Panel (Dros-OSP). Using this community resource, we collected transcriptomic and genomic data for 1800 individual flies and were able to map *cis* and *trans* eQTLs for 98% of the genes expressed in *D. melanogaster, increasi*ng by thousands the number of genes for which regulatory loci are known in this species. We described, for the first time in the context of an outbred population, the properties of local and distal regulation of gene expression in terms of genetic diversity, heritability, connectivity, and pleiotropy. We uncovered that, contrary to long-standing assumptions, a significant part of gene co-expression networks is organized in a non-modular fashion. These results bring the fruit fly to the level of understanding that was only available for a few other organisms, and offer a new mapping resource that will expand the possibilities currently available to the *Drosophila* community. This data is available at Drosophilaeqtl.org.

## INTRODUCTION

Two decades of genome wide association studies and QTL mapping experiments in a wide array of species have revealed that most polymorphisms associated with variation in complex traits map predominantly to non-coding regions, and are likely to be involved in gene regulation (Albert & Kruglyak, 2015; Alsheikh et al., 2022). This poses major challenges for the characterization of the functional consequences of sequence variation, and ultimately for the understanding of how genetic polymorphisms drive phenotypic variation. To address this problem, considerable effort has been dedicated to mapping genetic variants that regulate variation in gene expression levels genome-wide (i.e. expression Quantitative Traits Loci or eQTLs). While eQTL studies in humans have led the way (Stranger et al., 2007; The GTEx Consortium, 2020; Võsa et al., 2021), large-scale projects in model systems such as yeast (Albert et al., 2018; Kita et al., 2017), mice (Gonzales et al., 2018) and *Arabidopsis* (Lan et al., 2021) have, for each of these species, provided detailed annotations of their regulatory genome. These eQTL catalogs have proven invaluable resources that have enabled the identification of general characteristics of the genetic basis of gene expression variation across species. Finally, such catalogs have helped prioritize candidate genes by suggesting mechanisms through which polymorphisms impact gene function in the context of specific traits, diseases, or adaptations. Gene expression variation is regulated by a large number of genetic variants that can only be detected in experiments with high statistical power (Albert et al., 2018; Võsa et al., 2021). As a result, for most organisms, the gene regulatory landscape that explains individual transcriptional variation remains poorly understood. This is also true for *Drosophila melanogaster, which, d*espite having one of the best annotated genomes and being an iconic model system, has lagged behind in the identification of eQTLs regulating transcriptional variation at the population-level, limiting the reach of this powerful model system. Studies aimed at understanding the genetic basis of transcriptional variation in *D. melanogaster have, thu*s far, been limited to a relatively small number of inbred lines (Everett et al., 2020) or pools of F1 individuals derived from crosses of inbred lines (King et al., 2014), limiting the statistical power. In addition, the use of inbred genomes complicates the study of gene interactions, such as dominance or epistasis, which are an important part of the genetic architecture of most complex traits (Mackay, 2014).

To address these limitations, we created a new synthetic outbred mapping resource that can be used to dissect the genetic basis of any complex trait. We created five mapping populations, one for each geographic location represented in the wild-derived Global Diversity panel (Grenier et al., 2015) with flies collected in Ithaca (USA), the Netherlands, Beijing (China), Zimbabwe and Tanzania. For each location, 15 to 18 inbred lines were crossed using a round-robin design, and the resulting population was maintained in large cages and allowed to recombine freely for over ∼125 generations. The resulting *Drosophila Outbred Synthetic Populations* (Dros-OSP) differ from other mapping panels like DSPR (King et al., 2012) and DGRP (Mackay et al., 2012) in that individuals are not maintained as inbred lines. The key benefits of this design are: (1) It assures that the allele frequency spectrum is biased towards common alleles, in contrast with wild populations, which improves statistical power to detect associations with alleles that otherwise would be too rare in the wild. (2) More than ∼125 generations of random mating and recombination erode long LD blocks, which improves mapping resolution. (3) The outbred nature of the Dros-OSP allows for the exploration of complex traits in a natural genomic context, where complex interactions like dominance and epistasis do occur. (4) Finally, this mapping panel does not impose any limitations on sample size, experiments can be as large as necessary, which is particularly important for the identification of genotype-phenotype association with small effects sizes, as may be expected for most complex traits.

Using the *Drosophila* Outbred Synthetic Population from the Netherlands, we simultaneously collected DNA and RNAseq data for 1879 outbred flies across two tissues. We identified eQTLs for 98% of genes, increasing by thousands the number of genes for which regulatory loci are known in *Drosophila*. Such a comprehensive genome-wide description of the regulatory landscape allowed us to describe the properties of local (cis) and distant (trans) eQTLs in terms of genetic diversity, heritability, connectivity, and pleiotropy. Additionally, we use state-of-the-art clustering algorithms to uncover communities in the gene co-expression network that do not follow the traditional expectation of modularity maximization. This resulted in new insights into the organization of *Drosophila* transcriptional networks. The results of this study are available at *Drosophilaeqtl*.*org*, a website that provides a user-friendly interface to explore the properties of the *Drosophila* transcriptome organization and eQTL map.

## RESULTS

### A new mapping resource: The *Drosophila* Outbred Synthetic Populations (Dros-OSP)

Our ability to dissect the genetic basis of complex traits depends on the available mapping populations. Traditionally, inbred and recombinant inbred lines panels (RILs) have been a commonly used resource for complex trait mapping. However, research in mice and rats has shown that using outbred populations greatly improves mapping resolution and power to detect genotype-phenotype associations, while maintaining a genetic context closer to wild populations (Gileta et al., 2022; Pallares et al., 2015; Yalcin et al., 2010). Here, we present a new outbred mapping resource in D. *melanogaster (Dros-OSP*) for mapping complex trait variation.

The Dros-OSP represents a significant shift from existing mapping resources in *Drosophila*. Traditionally, the sequence of a given set of inbred lines (or RILs) is provided as a resource. This means that the user only needs to phenotype their trait of interest in the desired lines, and can benefit from the available genetic information to perform genetic mapping. This paradigm was a game-changer for the field when DNA sequencing was cost prohibitive. However, the limited number of available inbred lines imposes a cap on the statistical power to detect genotype-phenotype associations. And the fact that such associations tend to have small effect sizes for most complex traits, further limits the inferences made with small sample sizes. The availability of outbred mapping populations not only remedies the sample size limitations, improving statistical power, but also allows us to dissect complex traits in a population that is more similar to a natural population in that each individual has a unique heterogeneous genome. The trade-off is that the user will need to sequence every Dros-OSP individual used for mapping. However, novel methods and reduction of sequencing costs have made possible the high-throughput and low-cost collection of genomic and transcriptomic data in thousands of individuals (e.g., (Pallares et al., 2020)).

The *Drosophila* Outbred Synthetic Panel has ideal properties for genetic mapping. It has an allele frequency spectrum biased toward common alleles (Fig. S1A), harbors ample genetic diversity (1,128,092 segregating SNPs MAF>1%, Fig. S1B, D-I), has rapid LD decay (∼200bp, Fig. S1C), and shows virtually no genetic structure (the first SNP-based PC explains less than 3% of genetic variation, Fig. S2). And finally, the Dros-OSP populations represent genetic and phenotypic diversity of five locations around the world: Tasmania, Beijing, the Netherlands, Zimbabwe, and Ithaca (USA).

Here, we illustrate one application of this mapping resource to characterize the genetic architecture of transcriptional variation. Using the Dros-OSP population from the Netherlands, we have quantified genetic and transcriptional variation genome-wide in 1879 samples of outbred *D. melanogaster. We explo*red gene expression variation in two body parts, the head and the body, referred to as tissues thereafter.

We found that 9473 genes are expressed in adult female D. *melanogaster across bo*th tissues. Most genes are expressed in the two tissues explored here, head and body (n = 7795), but each tissue has ∼10% of genes that are reliably detected only in one tissue (tissue-specific), 1082 genes (12.6%) and 596 genes (10.4%) in head and body transcriptomes, respectively (Table S1). Overall, the expression level of genes expressed in both tissues (shared) is strongly correlated, although there is notable variation at the gene-specific level (Fig. 1A). This is reflected in the fact that 94% of these genes are differentially expressed between tissues (Fig. S3A, Table S1). The top differentially expressed genes are enriched in synapsis and sensory-related processes mediating locomotion and mating behavior (Fig. S3B). And, as may be expected, tissue-specific genes are enriched for visual perception and neuropeptide signaling in the head, and egg production in the body (Fig. 1B). Notably, tissue-specific genes are expressed at lower levels than shared genes (Fig. 1A).

**Figure 1.**
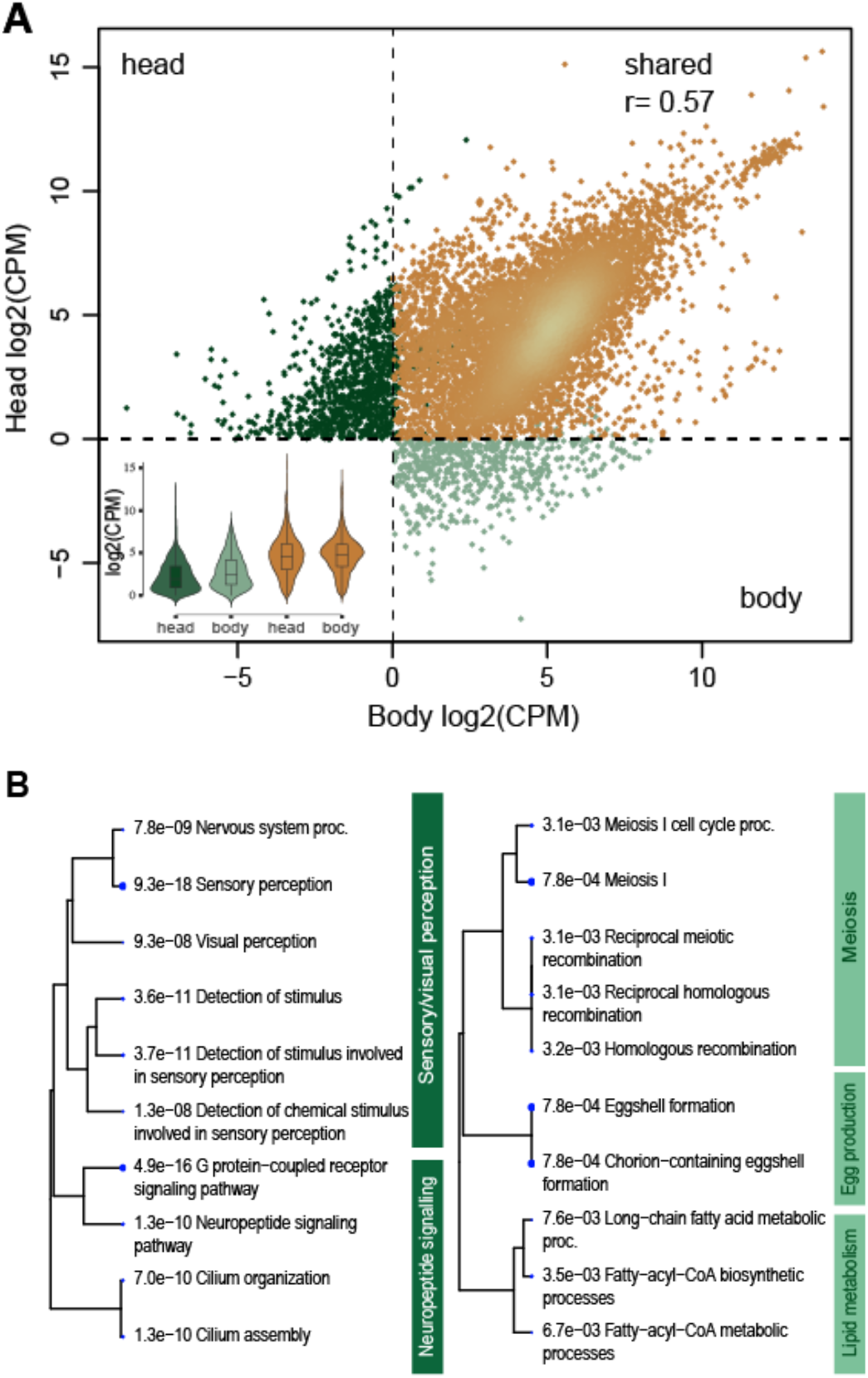
Correlation between head and body gene expression levels in D. *melanogaster transcrip*tomes. **(A)** Correlation between expression levels in head (940 samples) and body (939 samples). Each dot represents a gene; dotted lines indicate the threshold used to determine whether a gene is expressed or not (average CPM>1 and detected in at least 20% of the samples). Orange, 7795 genes expressed in both tissues. The Pearson correlation r, was calculated only including shared genes, p-value <2.2e-16. Dark green, 1082 genes with head-specific expression. Light green, 596 genes with body-specific expression. The inset plot shows the mean expression for tissue-specific (green) and shared genes (orange). **(B)** GO Biological Process enrichment for tissue-specific genes (dark green, head-specific; light green, body-specific). The top ten significantly enriched terms at FDR 0.05 are shown grouped by the number of shared genes between terms. Dot size indicates the p-value of the enrichment, which is also indicated for each term. Analysis done in ShinyGO v0.77 (Ge et al., 2020).

## Heritability of gene expression

We estimated SNP heritability per gene using the linear mixed model (LMM) implemented in GEMMA (X. Zhou & Stephens, 2012). This estimate corresponds to the amount of gene expression variance in the fly population explained by genetics effects. Non-zero heritability estimates were obtained for 72% (body) and 80% (head) of genes (Fig. S4-S6, Table S2). The heritability distribution is skewed towards low values, with an average of 0.07-0.08 (Fig. 2A). Genes with higher heritability are expressed at higher levels (Fig. 2B) and have higher nucleotide diversity, π (Fig. 2C) than low heritability genes. High and low heritability genes are not randomly distributed in the genome, chrX and chr4 are significantly enriched in low heritability genes, while the other main chromosomal arms (chr2L, chr2R, and chr3R) are enriched for high heritability genes. This pattern matches the differences in genetic diversity between chromosomes (Fig. S1D-I). These groups of genes also differ in their biological function (GO Biological Processes). Low heritability genes are enriched for very general regulatory processes including protein and RNA metabolism in both tissues, and early development and morphogenesis in the body transcriptome (Fig. S7). On the other hand, high heritability genes are enriched in more specific processes like carbohydrate and fatty acid metabolism, and immune response and detoxification (Fig. S8).

**Figure 2.**
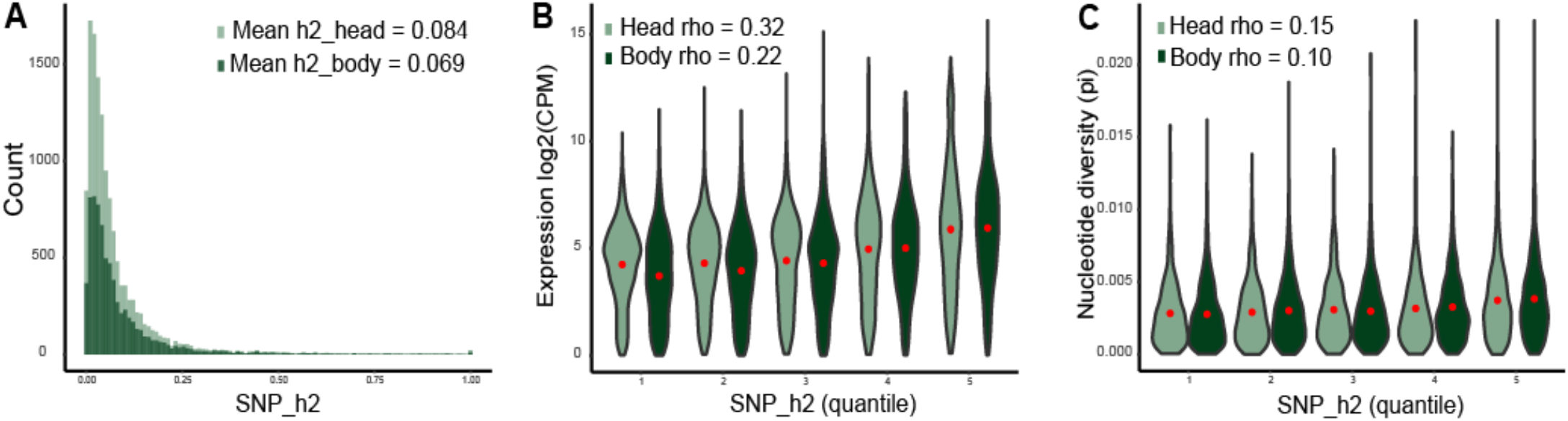
Heritability of gene expression. (A) The percentage of population-level expression variance explained by ∼380,000 SNPs (MAF>0.05), or SNP heritability (SNP_h2), is shown for 6001 genes expressed in the body, and 7155 genes expressed in the head. (B) There is a positive correlation between expression levels and SNP heritability. Genes in the first and fifth quantile have a median gene expression of 12.9 CPM and 63.6 CPM respectively, in the head (23.3 CPM and 57.5 CPM in the body). (C) There is a positive correlation between heritability and per-gene nucleotide diversity (π). Average per-gene π in first and fifth SNP heritability quantiles is 0.0027 and 0.0038, respectively in the head (0.0028 and 0.0037 in the body). The Spearman rank correlation is shown in B and C for each tissue, p-value for all tests < 2.2 e-16. Rank correlations were calculated using SNP_h2 as a continuous variable and not divided in quantiles.

### Structure of the *Drosophila* transcriptome

Next, we investigate patterns of co-expression to better understand community structure among transcripts. At the individual gene level, the fly transcriptome appears weakly connected. After removing poorly supported edges (Spearman correlation p-values > FDR 1%) the average correlation per gene is 0.13 in both tissues (min = 0.11, max = 0.24, Fig. 3A, Table S1), and the median number of genes connected to the focal gene (i.e., degree) is 329 and 469 in head and body transcriptomes, respectively (min = 3, max 4899, Fig. 3B, Table S1). However, although the average correlation per gene is rather weak, the correlation between specific gene pairs can be very high (max rho = 0.95, Fig. 3A).

**Figure 3.**
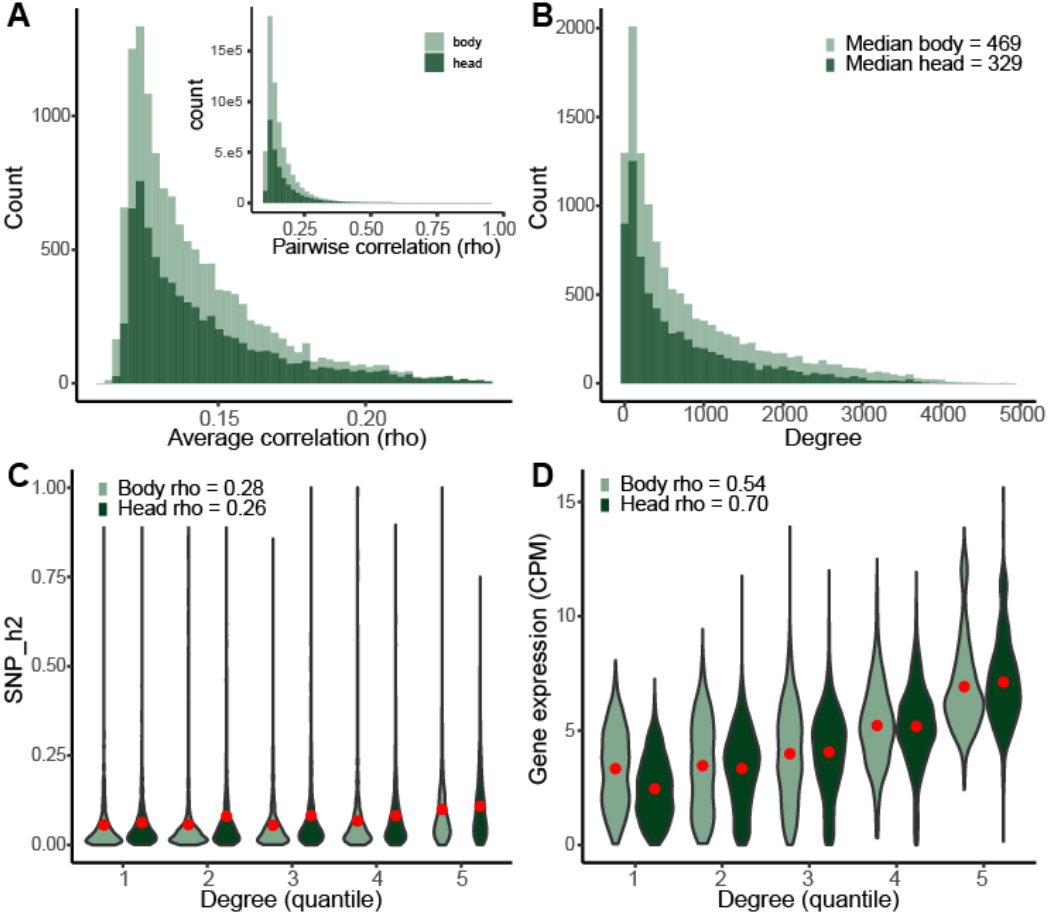
Patterns of gene connectivity based on expression correlation. Two gene-level connectivity measures are shown: (A) Average pairwise Spearman correlation between the focal gene and all other genes. Only correlations supported by a p-value below FDR 1% were used to estimate the average. Median correlation per gene is 0.13 in both tissues (min rho = 0.11, max rho = 0.24). The inset shows the distribution of pairwise correlation for all gene pairs with significant correlation p-values, highlighting the fact that although the distribution is skewed towards low correlations, some gene pairs are highly correlated (max rho = 0.95). (B) Degree represents the number of genes connected to a focal gene. For each gene, we counted the number of genes with significant Spearman correlation (FDR 1%) (min = 3, max = 4899). (C) Correlation between gene expression heritability and degree. Median SNP_h2 first vs fifth degree quantile: 0.034 vs 0.082 in the head; 0.026 vs 0.065 in the body (D) Correlation between gene expression levels (CPM) and gene degree. Median CPM first vs fifth degree quantile: 4.9 vs 111.9 in the head; 13.5 vs 79.5 in the body. Rank correlations in C and D were calculated using degree as a continuous variable, p-values for the correlations were both < 2.2 e-16.

Highly connected genes (correlated with ∼20% of genes) are enriched in ATP metabolism, and translation in both tissues, and also in behavior and synapsis in the head (Fig. S10). While genes with low degree (correlated with <1% of genes) are involved in RNA processing including ncRNA and tRNA, with specific modules in the head involved in DNA repair, and in carbohydrate and lipid related processes in the body (Fig. S11). Expression level, first, and SNP heritability, second, are the gene properties best correlated with connectivity metrics. Highly expressed, highly heritable genes are connected to a large number of genes, and such connections are supported by high correlation values (Fig. 3C-D, Fig. S9).

To go beyond the gene-level description presented above and explore the community structure of the D. *melanogaster transcrip*tome, we clustered genes using the Weighted Nested Degree Corrected Stochastic Block Model (SBM) (Karrer & Newman, 2011; Peixoto, 2017). While traditional clustering approaches try to maximize intra-modular correlation while minimizing correlation between modules, SBM uses a Bayesian approach to define gene clusters that does not rely on modularity (see Methods for more details on SBM, Fig. S12). Consequently, here we refer to clusters of co-expressed genes as “gene blocks” instead of “gene modules” to highlight the difference between these two approaches, and the different biological information that they capture. We further explore this approach in a separate manuscript (Melo, et al 2023) and briefly summarize the results below.

Body and head transcriptomes were partitioned into a total of 78 and 82 blocks, respectively, with a median of 32 and 36 genes per module (Fig. 4A, Table S3, Table S1). Such fine clustering of genes allows for a clear biological interpretation of most modules which are, on average, enriched in only six GO Biological Process terms (Table S3-5). Although the number of total modules, and the gene-level patterns described in the section above are very similar between head and body, there is a fundamental difference in the way the transcriptome is structured: the head transcriptome is highly modular, with 94% of the blocks resembling traditional co-expressed modules (i.e., higher within than between-block correlation, shown as positive assortativity values in Table S3), while only about ∼69% of the blocks that constitute the body transcriptome are modular (Fisher exact test, p-value 5.88e-5, OR 6.76) (Fig. 4B). This highlights the importance of not relying only on modularity maximization methods when studying transcriptome structure. Our clustering approach was able to recover biologically relevant information from thousands of genes that would have been missed with modularity-maximization approaches. A highlight of the results is that the same proportion of non-modular blocks has significant GO terms enrichment compared to modular blocks (60-70% of blocks have GO enrichment, Fisher exact test: p-value body = 0.59, p-value head = 1) (Fig. 4B).

**Figure 4.**
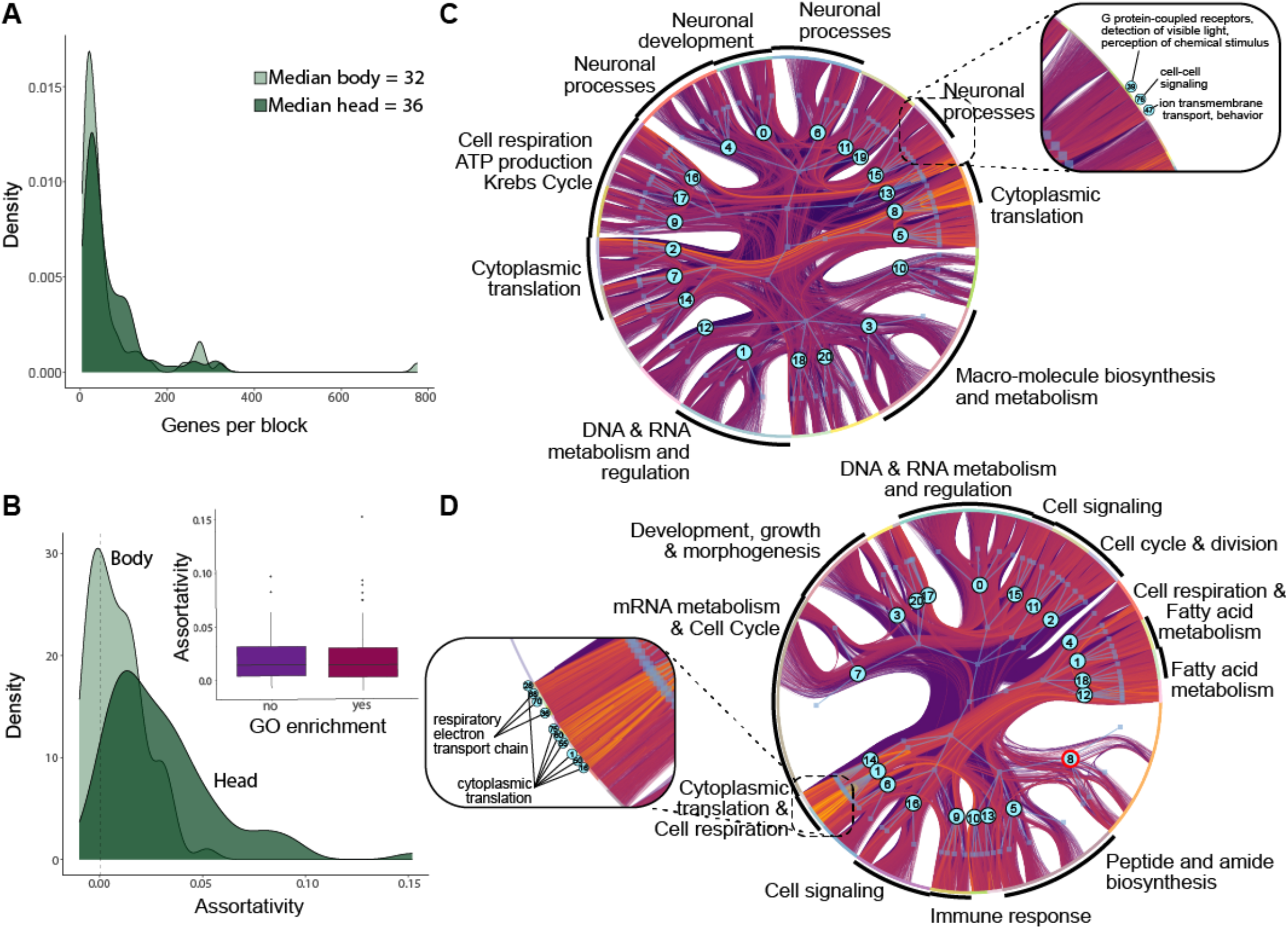
Modular organization of the fly transcriptome. The Drosophila transcriptome is highly structured, with 78 gene blocks in the body and 82 in the head. (A) The distribution of the number of genes per gene block is heavily biased towards small blocks (median = 34 genes), increasing the resolution of the GO enrichment for each block. (B) Not all gene clusters follow modularity assumptions. For each gene block, a level of assortativity was defined; values > 0 indicate modular behavior (i.e., genes within a block are more correlated with themselves than with genes in other modules). 94% of head blocks are modular, while only 67% are modular in the body. The inset plot shows that the level of assortativity is not correlated with GO enrichment, indicating that modular and non-modular blocks are associated with functional information (60-70% of blocks have GO enrichment, Fisher exact test: p-value body = 0.59, p-value head = 1). (C-D) Hierarchical representation of the transcriptome structure for head (C) and body (D). Light blue lines indicate the order in which the transcriptome was consecutively split into blocks (squares. The block in the middle of the circus plot represents the whole transcriptome. The blocks along the edge correspond to the final level of partition (level 1), that is, the smallest gene clusters whose characteristics are plotted in (A-B). GO enrichment terms for some level 2 blocks (numbered blue circles) are annotated. The numbered circles in the zoom-in panels indicate block ID in the last level of partition (level 1) and GO enrichment terms for those small blocks are annotated. 60% of gene blocks in the head and 70% are enriched for GO Biological Process terms regardless of the modular nature of the block. The zoom-in panels show the fine scale GO partitioning of an assortative block in the head (C), and a non-assortative block in the body (D).

### Genetic regulation of transcriptional variation

Consistent with the fact that most genes have non-zero heritability estimates, we identified eQTLs (genetic variants regulating expression levels) for 98% of the genes in the D. *melanogaster transcrip*tome, making this one the most comprehensive eQTL map in this species (Fig. 5, Table S2). For genes with eQTLs, 60% have at least one *cis-*eQTL (SNP position within ±10Kb of gene body), and 99% at least one *trans-*eQTL (Fig. S13, Table S1). And, 58% of genes have significant eQTLs in cis and trans, highlighting the complexity of gene expression regulation. A single gene has a median number of four *cis-*eQTLs and five *trans-*eQTLs. To further characterize the *cis-*regulatory landscape, we annotated all 258,283 *cis-*SNPs (SNPs located ±10Kb from any gene in the genome) based on the predicted effect that each allele will have on gene expression levels. To that end, we used DeepArk, a machine learning-based approach that uses 1552 experimentally derived *D. melanogaster datasets* to predict the cis regulatory activity of individual SNPs based on sequence information (Cofer et al., 2021). Compared to non-significant SNPs, the *cis-*eQTLs identified in this study are enriched for non-zero predicted regulatory effects, and particularly strongly enriched for large predicted effects (Figure 6A). This relationship between statistical discovery and predicted effect size based on experimental data has been previously shown for *cis-*eQTLs detected in genome-wide-association studies in humans (J. Zhou et al., 2018).

**Figure 5.**
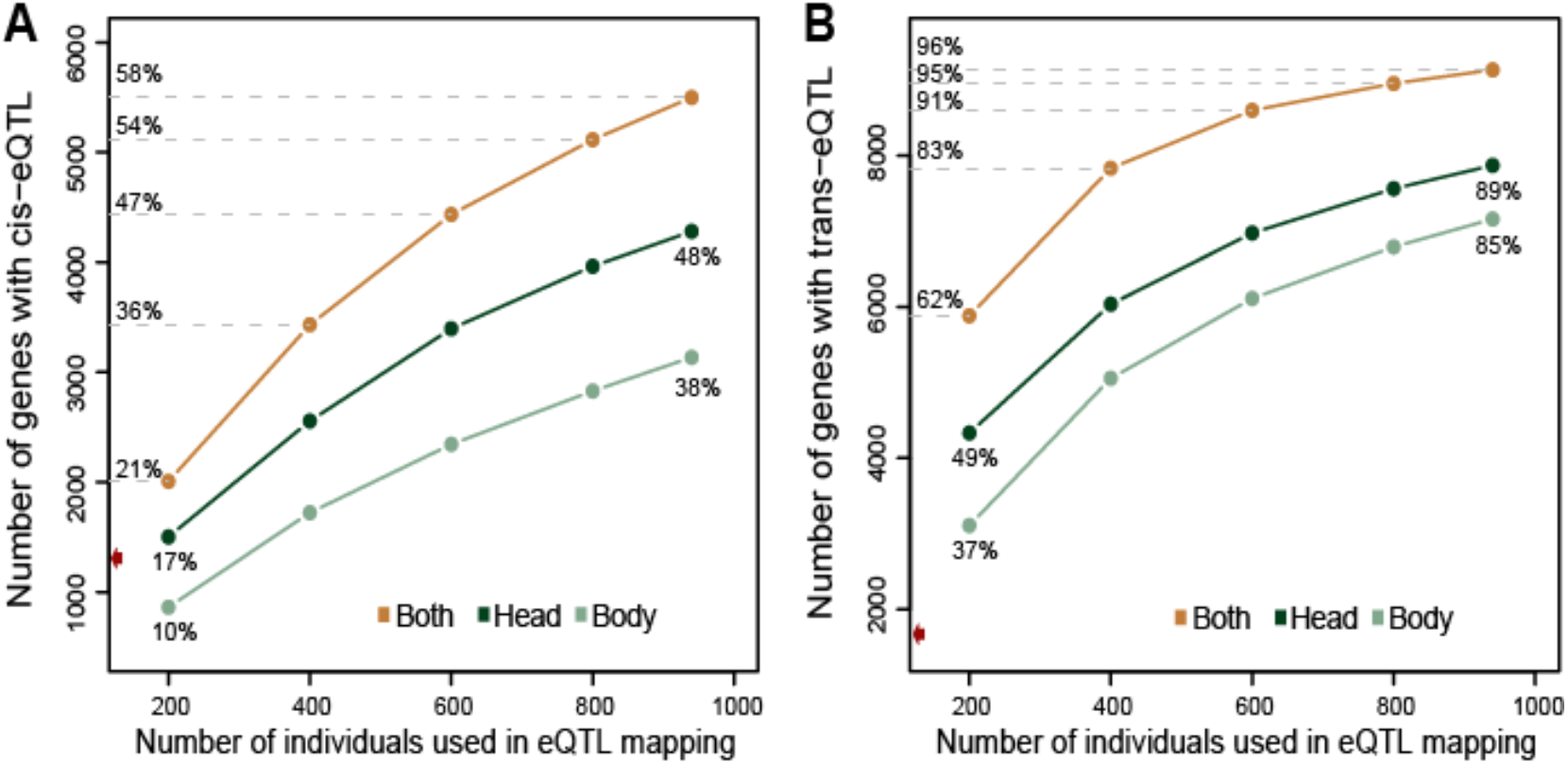
Saturation level of the eQTL map in D. *melanogaster*. *To evalu*ate the effect of sample size (i.e., statistical power) in the detection of (A) cis-eQTL and (B) trans-eQTL, the full dataset of ∼1000 flies per tissue was randomly down-sampled to smaller datasets in increments of ∼200 individuals per tissue (x-axis). To get the overview of the total number of genes with eQTL, the results of both tissues were combined (orange line). We identify cis-eQTL for 58% of the genes expressed in female D. *melanogaster (n = 5417*), and trans-eQTL for 97% of the genes (n = 9001). The red arrow on the y-axis represents the number of genes with eQTL from previous eQTL mappings in D. *melanogaster (1284 gen*es with cis-regulation, and 1653 with trans-regulation, (Everett et al., 2020).

**Figure 6.**
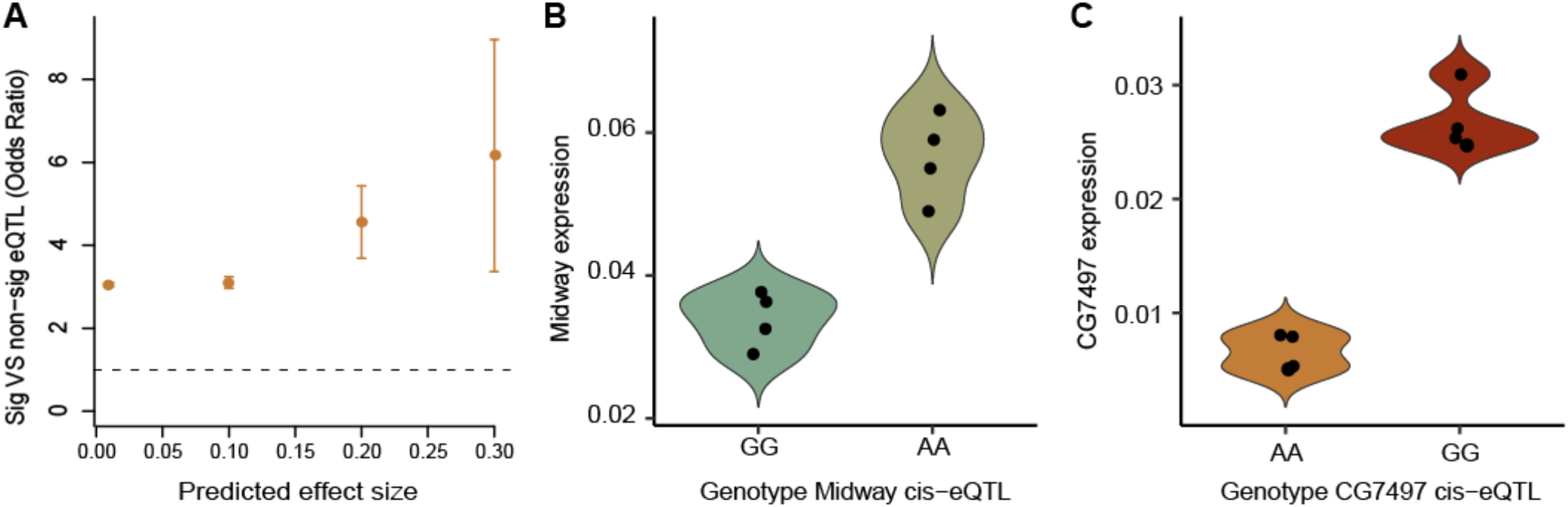
cis-eQTLs are enriched for large predicted regulatory effects and experimentally replicated. (A) The cis-eQTLs identified in this study using a statistical approached are enriched in large predicted regulatory effects, when compared to non-significant SNPs. Effect size prediction was done using the deep-learning algorithm implemented in DeepArk. Sample size for the comparison: 68,859 candidate cis-eQTL; 258,283 non-significant sites located in cis (within 10Kb from start or end of gene body). For each effect size, groups were compared using Fisher exact tests. All tests have p-values < 0.001. 95% confidence intervals for odds ratio are shown. (B,C) The genotype at candidate cis-eQTLs regulates expression levels of the focal gene. Each dot corresponds to a pool of 10 outbred flies fixed for one eQTL allele but with heterogeneous genomic backgrounds. Expression levels of the genes (B) midway and (C) CG7497 were quantified using rt-qPCR (midway t.test p-value: 0.0015, fold change 1.68x; CG7497 t.test p-value 8.088e-5, fold change 4.08x).

Our results show that such relationship is also true for the statistical inference of eQTLs in *Drosophila*. Finally, *cis-* eQTLs that have regulatory effects in both, head and body, tend to have larger predicted effect sizes compared to tissue-specific *cis-*eQTLs (Fig. S14A). To facilitate follow-up analysis, for each gene we annotate the best *cis-*eQTL as the one with the largest effect size predicted by DeepArk (Table S6A,B). With this, we offer an avenue for prioritizing candidate *cis-*eQTL for 58% (n=5499) of the genes expressed in female D. *melanogaster*.

*The ge*netic regulation of gene expression tends to be ubiquitous when in trans and tissue-specific when in cis. To allow this comparison, we focus on genes expressed in both tissues (n = 7719) and find that while 77% of the genes with trans regulation have *trans-*eQTLs in both tissues, only 41% of genes with *cis-*regulation have *cis-*eQTL in head and body.

As seen before in humans and other organisms, most eQTLs are located in non-coding regions of the genome (95% of *cis-* eQTL, 97% of *trans-*eQTL in *Drosophila*, Fig. S15B, Table S6). But, in contrast with humans, due to the smaller genome size in *Drosophila*, the majority of eQTLs fall in genic, although non-coding, regions (73% of *cis-*eQTL, 67% of *trans-* eQTL, Fig. S15B). Although this highlights the already well-established importance of non-coding genetic variation, we also find an important role for coding variation in regulating gene expression: eQTLs are enriched for SNPs that generate non-synonymous protein substitutions, compared to SNPs that are not eQTLs (chi-square test, *cis-*eQTL p<2.2 e-16, odds ratio = 2.18; *trans-*eQTL p<2.2e-16, odds ratio = 1.88). This result suggests two things: first, *trans-*regulation of gene expression can be mediated by protein changes in the focal gene where the eQTL lays (e.g., protein changes in a transcription factor will affect its binding to the promoter of distant genes). Second, and more interesting, genetic variation that results in protein changes in the focal gene also tends to modify its own expression levels (*cis-*eQTL). The importance of the latter mechanism is further supported by the fact that *cis-*eQTLs located within the gene region (as opposed to upstream or downstream) tend to fall in exons more than *trans-*eQTL, which instead are mostly found in introns (Fig. S15A).

We find that each eQTL is associated with the regulation of one or very few genes (median = 1). However, some eQTLs have tens to hundreds of *trans-*effects (max eQTL pleiotropy in head tissue = 56, in body tissue = 776, Table S6). This pattern suggests the presence of trans hot-spots. We identify 22 and 189 hot-eQTLs in head and body, respectively, defined as SNPs regulating more than 20 genes in trans. The pattern of hotspots differs between body parts, with the body transcriptome showing more hotspots, but also more of them seem to be clustered around telomeric regions which suggest that they might be driven by strong linkage in those regions (Fig. S17). These highly pleiotropic hot-eQTLs represent the full spectrum of allele frequencies in the fly population (Fig. S16B), and although preferentially located in pericentromeric regions, hot-eQTLs are found across the genome (Fig. S16C,D). These highly pleiotropic SNPs are necessarily regulating genes in trans, but to infer whether such effects might be direct (the eQTL regulates distant genes through e.g., long-range chromatin interactions) or indirect (the eQTL regulates expression of a gene in cis, and the gene product regulates the expression of distant genes), we asked whether hot-eQTLs are more likely to regulate at least one gene in cis (SNP is both, *cis-*eQTL and *trans-*eQTL) or to only regulate distant genes (SNP is a *trans-*eQTL but not a *cis-*eQTL). We find that for most hot-eQTLs an indirect mechanism is more likely because they are more likely than not to be a *cis-*eQTL (odds ratio 1.9, p-value Fisher test = <2.2e-16, Fig. S16A, Table S7).

The experimental validation of regulatory effects on gene expression currently presents a challenge, given that large population sizes are needed to detect the small effects associated with eQTLs. Here, we have used a simple experimental design that relies on pools of individuals that share the same genotype at the focal *cis-*eQTL while retaining heterogeneity in their genomic background (Wolf et al., 2023). This approach reduces expression variation within a genotype while increasing the statistical power to detect differences in mean expression between alternative genotypes. Using this approach, we show one of the very few validations of the regulatory effect of naturally segregating genetic variation using outbred individuals (Fig. 6B, C, Fig. S14B-D).

## DISCUSSION

To understand how the transcriptome is structured and regulated at the population level, we used the *Drosophila* Outbred Synthetic Population derived from the Netherlands. Using a total of 1879 genomic and RNAseq samples derived from two body parts, we identified genomic loci (eQTL) regulating variation in gene expression for 98% of the genes expressed in female D. *melanogaster. This com*prehensive genome-wide eQTL map enabled us to dissect general properties of cis and trans regulation in this species, bringing D. *melanogaster to a leve*l of understanding previously available only for a few species such as yeast (Albert et al., 2018) and humans (The GTEx Consortium, 2020).

Focusing on an outbred population, where each individual fly has a unique genomic composition, allowed us to collect a large-enough sample size to reach saturation of the eQTL map at the gene level, that is, 98% of the genes have at least one eQTL. Our eQTL catalog offers a comprehensive resource for the complex trait community, where the number of genes with known *cis-*regulation increased from 1284 (Everett et al., 2020) to 5417, and from 1653 (Everett et al., 2020) to 9001 for *trans-*regulation. In addition to the obvious increase in power derived from a large sample size, the use of two tissues allowed us to detect even more eQTLs. This is reflected in the fact that 59% of genes with *cis-* regulation and 23% of genes with *trans-*regulation have tissue-specific *cis-* and *trans-*eQTL, respectively. This result suggests that to further improve the eQTL map, in particular the *cis-*eQTL map, more tissues rather than sample size, are necessary.

We find that the expression level of most genes is regulated by both local and distal genetic variants, and that multiple SNPs are usually involved. Such pattern has been previously observed in humans, mice, yeast and fruit flies (Albert et al., 2018; Gonzales et al., 2018; Huang et al., 2015; Jansen et al., 2017; The GTEx Consortium, 2020). Despite such idiosyncratic gene-specific regulation, we find transcriptional regulation hotspots, where one single eQTL is associated with expression variation of tens or hundreds of genes. Although some hotspots can be attributed to batch effects or population structure (Kang et al., 2008), they seem to be a property of the eQTL map of several species, including mice (Gonzales et al., 2018; Hasin-Brumshtein et al., 2016), *Drosophila* (King et al., 2014), and yeast -where hotspots have been experimentally validated (e. g. (Albert et al., 2018; Smith & Kruglyak, 2008; Zhu et al., 2008)).

Recent machine learning approaches use sequence-based information to predict the impact of individual alleles on expression levels (Cofer et al., 2021; J. Zhou et al., 2018). However, they cannot predict which gene will be regulated by certain SNP. In species with small genomes and high gene density like D. *melanogaster, single S*NPs might act as *cis-* eQTL for many genes. On the other hand, statistical approaches like the one used here to map eQTLs, offer a direct link between a genetic variant and its target gene. To make use of these two powerful lines of evidence, we have annotated thousands of *cis-*SNPs (located ± 10Kb from any gene) with the statistical probability that they regulate a specific gene, and the machine learning-derived predicted effect size of such regulatory effect. This resource will help to prioritize SNPs for further experiments, as we have done here to validate multiple candidate *cis-*eQTL (Table S8). For such validation we implemented an approach that relies on using outbred flies fixed for a candidate eQTL allele while preserving their genomic heterogeneity (Wolf et al., 2023). To our knowledge, this is the first time that eQTL effects have been experimentally validated in outbred individuals. Traditionally, our understanding of the organization of transcriptional networks has been driven by the concept of modularity, where a module is defined as a group of genes that are more strongly connected to each other than to genes outside that module. Here, we implemented a method that does not rely on modularity maximization to define gene clusters, the Stochastic Block Model – SBM (Karrer & Newman, 2011; Peixoto, 2017). Doing so, we discovered that a considerable proportion of gene blocks in *Drosophila* do not show the expected pattern of high connectivity within modules and low connectivity across modules. This suggests that a full understanding of transcriptional organization needs to go beyond modularity and incorporate other properties of gene connectivity. For example, two tightly correlated groups of genes can be clustered in separate blocks because the genes in the first group are correlated to a third block while the genes in the second block are not. It’s also possible that a group of genes is clustered in the same block not because they are correlated among themselves but because they tend to be correlated with the same genes in another block. Elsewhere we discuss in detail the advantages of using methods that don’t rely on modularity maximization (e.g. SBM) to study the architecture of transcriptional networks (Melo et al., 2023).

This study uncovers the fine scale landscape of gene expression regulation in D. *melanogaster, identifi*es general properties of cis and trans eQTL, and describes the global organization of the transcriptome. In doing so, it sets the ground for future studies using gene expression as means to understand how genetic variation gets translated into phenotypic variation. Critically, defining the direction in which information flows in transcriptional networks by building directed transcriptional networks will allow us to draw the causal path from allele frequency to phenotypic variation.

## ACKNOWLEDGEMENTS

We specially thank Paul Durst, Amanda Lea, Luke Henry, Michael Fernandez, Minjia Tang, Annett Schmittfull, and Sudarshan Chari for help during the two weeks of daily fly collection. And Annett Schmittfull, Serge Picard, and Elena Filippova for help processing the samples. Thanks to Jen Grenier and Andy Clark for sharing the VCF file for GDL. Thanks to Lance Parsons for bioinformatics support and to Andy Dahl for discussions on the statistical approach to map eQTL in this population. During this experiment, our incubator broke down carbonizing all flies at 40 °C; thanks to the backup that Andy Clark and Jen Grenier kept at Cornell, we were able to repeat the experiment.

## DATA ACCESS

Supplementary tables S1-S5, S7 and S8 are in the Supplementary Tables excel file. Supplementary tables S6A-D (folder SupplementaryTables), files needed to replicate the eQTL mapping (folder GEMMA), raw gene count tables (folder GeneCountMatrices), and VCF files (folder VCF) can be found in: https://edmond.mpdl.mpg.de/privateurl.xhtml?token=c2fc74ee-accd-4b2a-82f3-0829ecf5951b (this dataset is currently privately shared and will be made public upon publication). In the webpage [*Drosophilaeqtl*.org](http://Drosophilaeqtl.org) the results of this study can be explored at the level of SNP, gene, and block.

## AUTHOR CONTRIBUTIONS

Conceptualization and design were by L.F.P. and J.F.A. DNA and RNAseq data generation were by L.F.P, J.P and V.A. Data analysis by L.F.P, D.M. and E.C. Website design was by S.W. Funding from JFA. Manuscript writing was by L.F.P., D.M. and J.F.A., with input from all authors.

## MATERIALS AND METHODS

### A synthetic global population of *Drosophila melanogaster*

*Using the* Global Diversity Lines (Grenier et al., 2015) as founder lines, we created five *Drosophila* Outbred Populations that represent genetic diversity from five continents – Tasmania, Beijing, The Netherlands,Zimbabwe, and Ithaca (USA). For each geographical location, 14 to 20 Global Diversity Lines were crossed in a round-robin design. The resulting F1 flies were again crossed in the same way to avoid losing any of the parental chromosomes. The F2 individuals were combined in one cage per geographical location and have been freely recombining for ∼200 generations at a population size exceeding ∼5000 flies. Each individual fly in these populations has a unique genomic composition, making them suitable for mapping studies in a context analogous to a natural population. In this study, we used the outbred population from The Netherlands, which will be called Nex in the text. At the time of the experiment, flies had been recombining for ∼120 generations, with a population size in the order of several thousands. The population was slowly expanded in a large cage for ∼10 generations until reaching ∼10k individuals before the experiment was conducted. All flies were maintained at 25 °C, 65% humidity, and a 12h:12h light:dark cycle, and were fed media with the following composition: 1% agar, 8.3% glucose, 8.3% yeast, 0.41% phosphoric acid (7%), and 0.41% propionic acid (83.6%).

### Experimental design

The experimental flies were collected from the Nex outbred population in three consecutive rounds of egg laying from the same parents. Each egg lay consisted of 15 bottles of fresh media placed inside the cage for ∼24 hours or less, controlling as much as possible for equal egg density between bottles. Newly eclosed adult flies were transferred to new bottles at the same time every day to assure that all flies were age matched; flies were collected for about a week until most pupae had eclosed. After allowing one or two days for all flies to mate, males and females were sorted and placed in different bottles until they were seven days old. To avoid CO2 exposure, we worked in a 4°C room to cold-shock seven-day old females and plated them in 96-well plates that were immediately submerged in liquid nitrogen to flash freeze the flies. Individuals that originated from the same egg lay and eclosion batch were always kept in the same plate (allowing us to account for potential batch effect in downstream analysis). Heads and bodies were then separated in the following manner: each plate was sealed with a rubber lid (Thermo-Fisher, #AB0566) and placed in liquid nitrogen for a couple of minutes, then the plate was bashed against a solid surface causing the head to fall off while the thorax and abdomen remain together; using a custom-made sieve, heads only were transferred to a new pre-cooled 96-well plate while bodies remained in the original plate. Head and body plates were sealed and immediately placed at -80°C for storage. All samples used in this study are female flies.

### mRNA extraction and RNAseq library preparation

We used a previously optimized low-cost and high-throughput protocol for mRNA extraction from single fly heads (Pallares et al., 2020); the protocol is described in detail here: Suppl. File 2 of (Pallares et al., 2020). RNAseq library preparation was done following the TM3’seq protocol (Pallares et al., 2020); a detailed description of the protocol, primers, and reagents is available in Suppl. File 1 of Pallares et al. More details can be found in Supplementary Methods.

### Processing of RNAseq data

Low-quality bases and adapter sequences were trimmed from the raw RNAseq reads using Trimmomatic 0.32 (Bolger et al., 2014). Reads shorter than 20nt after trimming were discarded. [Trimmomatic parameters: SE ILLUMINACLIP:1:30:7 LEADING:3 TRAILING:3 SLIDINGWINDOW:4:15 MINLEN:20]. The trimmed reads were mapped to the *Drosophila melanogaster genome r6*.14 using STAR (Dobin et al., 2013), and uniquely mapped reads were assigned to the set of 17727 genes annotated for the r6.14 genome using feautureCounts from the package Subread [parameters: -t exon –g gene_id] (Liao et al., 2014). At this point, only RNA samples with more than 500,000 reads assigned to genes and matching DNA samples (see below) were kept for further analysis. Finally, only genes present on the autosomes (2L, 2R, 3L, 3R, 4) and chromosome X were kept for downstream analysis. This resulted in 940 samples with RNA derived from head tissue (average gene counts per sample: 3M), and 939 with RNA derived from body tissue (average gene counts: 2.4M) (Fig. S17). At the gene level, only genes with mean CPM>1 and we only kept genes for which at least 20% of individuals in the population were assayed (with CPM>1 for the particular gene). These filters removed not-expressed and lowly expressed genes which corresponded to roughly half of the 17727 genes present in the Dmel r6.14 annotation file (51% of the head genes, and 53% of body genes). The final dataset used for further analysis consists of RNA samples that have DNA matching samples (see section ‘Processing of DNAseq data and SNP calling’): 940 RNA samples derived from head tissue expressing 8877 genes, and 939 RNA samples derived from body tissue expressing 8391 genes. 593 samples were derived from head and body tissue from the same individual fly, while the rest (n=693) correspond to one tissue per fly. The vast majority of genes were detected in more than 90% of the samples (79% and 76% of head-and body-expressed genes). On average, each gene was detected in 873 and 858 samples in head and body tissue, respectively (median = 938, 931 respectively).

### DNA extraction and library preparation

DNA was extracted from the body or head lysate used to first extract mRNA in the following way: after mRNA bound to the Dynabeads™ in step 13 of the RNA extraction protocol (see https://lufpa.github.io/TM3Seq-Pipeline/mrna_extraction), the supernatant was transferred to a deep 96-well plate and mixed with 400ul of genomic lysis buffer (Zymo, #D3004-1). The mix was stored at -20C to await further processing. Samples were transferred to an Acroprep advance 1mL DNA binding plates (Pall Life Sciences, #8132) for DNA extraction using the Multi-Well Plate Vacuum Manifold (Pall Life Sciences, #5017) and DNA pre-wash, and gDNA wash buffers from Zymo (#D3004-1, #D3004-5). DNA library preparation was also performed in 96-well plates, following the tagmentation approach outlined in Picelli et al. (2014) and implemented in a CyBio® FeliX liquid handling robot (Analitik Jena). Details can be found in Supplementary Methods.

### Processing of DNAseq data and SNP calling

DNA reads were mapped to the D. *melanogaster reference* genome r6.14 using BWA (Li & Durbin, 2009) while preserving lane identity for samples that were sequenced in multiples lanes or batches. Secondary alignments were removed, and duplicated reads were marked using the function MarkDu-plicates in Picard (http://broadinstitute.github.io/picard). Samples with <500 PE reads were removed. The distribution of mean coverage per sample can be seen in Fig. S18 (average coverage of samples with head RNAseq data = 6x, and samples with body RNAseq data = 6.6x). To first create a set of high-quality SNPs for this population, we used a set of 210 samples with coverage > 8x (mean coverage 10x). Polymorphic sites were called for individual samples using HaplotypeCaller (-ERC GVCF), and joint variant calling for all samples was done with GenotypeGVCFs, both implemented in GATK v4.1.3.0. The resulting set of SNPs were hard-filtered following the Best Practices recommendations from GATK adjusted to the empirical distribution of our SNPs parameters. Using SelectVariants, we implemented filters at two levels, first to account for overall coverage parameters (remove if: singleton, total coverage <500 or >5000, average coverage <5x or >25x, present in <50% individuals), and second to account for site-specific quality parameters (remove if: QD<5.0, MQ<50.0, FS>10.0, SOR > 0.2, MQRankSum < -2.0 or > 2.0, -ReadPosRankSum < -2.0 or > 2.0, ExcessHet< 10.0 -this corresponds to a probability of violating HW equilibrium due to heterozygote excess < 0.1). These filters resulted in a high-quality set of 1,128,092 biallelic SNPs with MAF>1% that we refer to as dbNex. To verify the quality of the SNP calls in the Nex population used here, we compared dbNex with SNP datasets from DGRP2 (Huang et al., 2014; Mackay et al., 2012) and the Global Diversity Lines (Grenier et al., 2015); 87% and 97.6% of the dbNex sites were validated in these populations, respectively.

Using dbNex as “known dataset”, we performed base quality score recalibration (BQSR) for the 1286 samples that had matching head or body RNAseq data, and then we called SNPs using HaplotypeCaller and GenotypeGVCFs in GATK. We attempted extensively to optimize GATK’s VQSR algorithm to filter our final set of variants based on parameters learned from dbNex, DGRP2, and GDL SNP sets, but we did not obtain reliable thresholds to call high-quality SNPs. Therefore, we decided to filter the final set of SNPs using hard thresholds similar to the ones described above following GATK recommendations adapted to the empirical distribution of parameters from our fly samples (filters used in GATK’s SelectVariant; keep if: ExcessHet<54.69 – corresponds to a probability to violate HW due to heterozygote excess 3.4e-6, biallelic site, present in at least 500 individuals, mean coverage <22x, MQ>50, QD>2.0, FS<60, SOR<3, -2.0<MQRankSum<2.0, -2.0<ReadPosRankSum<2.0). Additionally, given that many of our individual samples have low coverage (see Fig. S18), we set to NA all individual genotypes with Genotype Quality (GQ) <9 corresponding to a level of confidence of 87.5%. To do this, we first mark these sites using VariantFiltration, and then SelectVariants --set-filtered-gt-to-nocall. 52 Individuals with <20% genotyping rate were removed. Finally, for each tissue, sites with MAF<5%, present in less than 20% of individuals, and with HW p-adj for heterozygote excess <1e-7 and heterozygote deficit <1e-40 were removed. We used two different thresholds for HW equilibrium because heterozygote excess is highly correlated with technical artifacts while heterozygote deficit is a known consequence of low-coverage datasets like ours (Benjelloun et al., 2019). These filters resulted in a set of 854,350 SNPs. Using Plink (Purcell et al., 2007), we removed one member of SNP pairs with r2>0.8 using a sliding window of 1000 SNPs with a 5 SNP offset. This step removed over half of the sites, resulting in 389,896 sites used for mapping. As a final quality check, we removed 4 samples which relatedness was larger than 1 using --rel-cutoff 0.99 in Plink. The final VCF file consists of 1286 samples matching 940 head RNA samples and 939 body RNA samples.

Coverage, genotyping rate per individual and number of individuals with genotype data for each site are shown in Fig. S18.

### Patterns of genetic variation in the mapping population

To characterize the genetic structure and variation in the mapping population we analyzed all 1286 DNA samples and 854,350 SNPs passing the quality control filters described above, but that had not been pruned based on LD. Using VCFtools (Danecek et al., 2011) we first estimated r2 between all pairs of SNPs in 1000bp windows (--geno-r2 --ld-window-bp 1000) excluding the pericentromeric and telomeric low recombination regions by removing the first 5Mb and last 15Mb from chr 2L, 2R, 3L, 3R and X, and the first 100kb and last 900kb of chr 4. The plots shown in Fig. S1 represent the smoothed relationship between LD and genomic distance calculated using Loess regression implemented in the loess function in R, using the smoothing parameter span=0.50. MAF was calculated using --freq in VCFtools; note that the allele frequency spectrum is truncated at MAF 0.05 because the sites used here were previously filtered based on MAF. Inbreeding coefficient (f) was estimated using the --het function, and relatedness using --relatedness. Nucleotide diversity per-site (π) was estimated in a subsample (n=320) of the best covered DNA samples (8x > coverage < 12x). VCF files for these 320 samples were re-generated to include variant as well as invariant sites using GenotypeGVCFs with the flag --include-non-variant-sites in GATK. Then, in VCFtools, while removing indels (--remove-indels) and keeping only mono-and bi-allelic sites (--min-alleles 1, --max-alleles 2), π per site and π in 1Kb windows was estimated using --site-pi and --window-pi 1000, respectively. Windowed π estimates were used to estimate chromosomal level π shown in Fig. S1, and using custom scripts π per gene was estimated as the average of per site-pi across gene length (see Fig. 2).

To evaluate whether large-scale genetic structure was present in the Nex population, we first generated the Genetic Relatedness Matrix (GRM) for all samples using SNPs pruned for LD>0.8 (n=389,896) using the command –make-grm-bin in GCTA (Yang et al., 2011), and then extracted the first 10 PCs with the option –pca (Fig. S2).

### RNAseq data normalization and covariates

Raw RNAseq counts per tissue were normalized by library size and composition using the function voom (Law et al., 2014) implemented in the R package limma (Ritchie et al., 2015) and the normalization factors estimated with calcNormFactors from the package edgeR (Robinson et al., 2010). The resulting data (log2(CPM)) is normally distributed, conforming to linear modelling assumptions and controlling for outliers typical of RNAseq count data.

All samples were processed in 96-well plates according to their respective round of egg lay and eclosion date (i.e., each plate contains samples from the same egg lay and eclosion date) to library sequencing, and therefore we use plate ID as the main variable accounting for batch effects in our dataset. For RNAseq samples derived from body tissue, accounting for plate effect was enough to remove latent structure in the data, and it was therefore used as the only covariate for downstream analysis. In contrast, in RNAseq data derived from head tissue, plate ID was not enough to remove batch structure and therefore we used the first three surrogate variables (SV) in addition to plate ID as covariates in follow-up analysis. SV discovery was done with the R package sva (Leek & Storey, 2007) using a null model that included plate ID as a predictor.

### eQTL mapping

To identify genetic variation regulating transcript abundance, we used GEMMA (X. Zhou & Stephens, 2012). Details on the comparison between GEMMA and Matrix eQTL (Shabalin, 2012) can be found in Supplementary Material and Fig. S19.

Given that not all individual flies were sampled for both, body-RNAseq and head-RNAseq (i.e., head and body tissue taken from the same individual), the individuals used for eQTL mapping in each tissue are different. We therefore estimated in GEMMA two GRM, one for each tissue, using the 389,896 quality-filtered and LD pruned SNPs (MAF>0.5) using the following parameters: -grm -miss 0.5 -gk 2. Voom-transformed gene counts were modelled independently for each tissue: head eQTL mapping, 940 samples, 8877 genes, plate ID and SV1-3 as covariates; body eQTL mapping, 939 samples, 8391 genes, plate ID as covariate. GEMMA parameters: -miss 0.5 -lmm 1.

To distinguish *cis-*eQTL and *trans-*eQTL, the results from GEMMA were processed in the following manner: First, cis SNPs were defined as sites within 10kb from the gene body, and trans SNPs as any other SNPs in the genome. Out of the 8877 genes expressed in the head, 8834 have SNPs in cis. Out of the 8391 genes expressed in the body, 8357 have SNPs in cis. This resulted in a total of 675,425 cis and 3,153,807,913 trans SNP-gene tests in the head dataset, and 632,149 cis and 3,096,239,321 trans tests in the body dataset. Then, because not all p-values were recorded due to the very large number of tests, p-values for *cis-* and *trans-*eQTL tests were adjusted for multiple testing using an FDR approach modified for the situation where p-values for all tests are not known, but the total number of tests is known. For this, we used the function p.adjust implemented in R (R Core Team, 2021), setting n to the known number of cis and trans tests. A threshold of 5% FDR was used to determine significance in each tissue. Shared eQTLs between tissues were defined as eQTLs with FDR < 0.05 in both tissues, and tissue-specific eQTLs as FDR < 0.05 in one tissue and FDR > 0.05 in the other.

### Heritability estimates

We estimated the amount of variance in gene expression explained by the set of 389,896 LD-pruned SNPs (MAF>0.05) and the standard error of the estimate using GEMMA (X. Zhou & Stephens, 2012). GEMMA provides the percent variance explained (PVE) which corresponds to the ‘SNP heritability’. The minimum SNP heritability output by GEMMA is 9.9999e-6 and is usually associated with extremely large error intervals. Given that there is no confidence in such estimates, we exclude those genes from further analyses involving SNP heritability estimates.

### Saturation curve

To assess the effect of sample size in eQTL discovery, we mapped eQTLs using subsets of 200, 400, 600 and 800 samples, in addition to the full dataset of 939 body and 940 head samples (Table S2). The datasets were created in the following way for each tissue: the full dataset of raw gene counts was randomly down-sampled to 200 individuals, genes with CPM<1 and detected in less than 20% of the samples were removed. The treatment of the raw RNAseq counts, eQTL mapping, and heritability estimates was the same as described before for the full dataset.

### SNP annotation

We annotated the SNPs used for eQTL mapping using two approaches. First, we used SNPeff (Cingolani et al., 2012) to determine the position in the genome (genic, intergenic, intron, exon, etc.) and the predicted impact on protein coding sequences (synonymous or non-synonymous changes) for all 389,896 SNPs. Secondly, we used the deep learning approach implemented in DeepArk (Cofer et al., 2021) to predict the regulatory impact of *cis-*SNPs (SNPs located within 10Kb of any gene) on the expression of their *cis-*genes (genes identified in our eQTL mapping). DeepArk uses 1552 D. *melanogaster experimen*tal datasets to predict regulatory impact based solely on the DNA sequence surrounding the SNP of interest. These datasets cover a wide range of tissues and developmental stages, but here we focus on the predictions based on the 245 assays performed in adult *Drosophila* because this is the developmental stage used in the current study. For each *cis-*eQTL we estimate the impact of a SNP on gene regulation as the predicted effect of reference allele minus predicted effect of alternative allele (ref-alt), and report the largest predicted effect size out of the 245 predictions made by DeepArk. This measure reflects the importance of a site in impacting gene regulation, i.e., if the predicted effect of reference and alternative alleles is very different, then the SNP potential to be involved in regulation of gene expression is high. If the effect, on the contrary, is very similar, then having one or the other allele is not relevant for determining the levels of expression of the focal *cis-*gene.

### Differential gene expression analysis

To detect the genes that are differentially expressed between head and body, raw gene counts of genes expressed in each tissue were merged, resulting in 7795 genes common to both tissues. The raw gene count dataset containing both tissues was processed through the following steps: first, TMM normalization factors were calculated with the calcNormFactors function from the R package edgeR (Robinson et al., 2010). Then, using the voom function in the R package limma (Law et al., 2014; Ritchie et al., 2015) raw counts were log-transformed and normalized using the normalization factors estimated in edgeR to control for library size and library composition, and precision weights for each gene-sample pair were estimated to control for the mean-variance relationship inherent to RNAseq data. Finally, given the presence of latent factors driving structure in the head transcriptomes (see section “RNAseq data normalization and covariates”), surrogate variables (SV) were estimated for the joint voom-transformed dataset using the R package sva (Leek & Storey, 2007). The first four SVs were sufficient to correct for the remaining structure in head transcriptomes and therefore were used as covariates, in addition to plate ID (see section “RNAseq data normalization and covariates”).

As described in the section “eQTL mapping”, some RNAseq samples were derived from the head and body of the same individual fly, making our dataset partially paired, with 801 independent samples (401 flies with only head RNA, and 400 flies with only body RNA) and 539 paired samples (539 flies with RNA data from head and body). To account for this, we estimated the within-individual correlation for each gene using the function duplicateCorrelation from the R package limma (Ritchie et al., 2015) on the voom-derived object, while specifying the model design and blocking the individual fly ID. Lastly, we modelled differential expression between tissues by fitting a linear model to the voom-derived object (contains log-transformed normalized counts and precision weights) using the function lmFit in the same R package, and including plate ID and SV1-SV4 as covariates. Parameters used in lmFit: block=individual ID, correlation=consensus correlation obtained from duplicateCorrelation as explained above. Differentially expressed genes were called at 5% FDR.

### Transcriptional structure and modularity

To explore the structure of D. *melanogaster transcrip*tome, we used a subset of the best covered samples per tissue, each one with more than three million RNAseq reads assigned to genes (average gene counts: head = 4.65M, body = 4.58M). This resulted in 248 body-RNAseq samples, and 318 head-RNAseq samples that were down-sampled to 248 to match the sample size of body transcriptomes. At the gene level, we kept the 5538 and 5269 genes expressed in body and head, respectively, that had mean CPM>1 and that were detected in all samples. Such filters result in a dataset that allows us to get more accurate estimates of the correlation strength between gene pairs, especially those that involve lowly expressed genes.

The raw gene count matrices were voom-transformed and surrogate variables were estimated for the head dataset as described in sections “eQTL mapping” and “Differential gene expression analysis”. To render gene count matrices ready for modularity analysis, we removed the effect of batch effects from the voom-transformed matrices (head batches: plate ID and SV1-2; body batches: plate ID) using the ‘function removeBatchEffect from the R package limma (Ritchie et al., 2015). We estimated the Spearman correlation for all gene pairs in the batch-free expression matrix, and only retained edges supported by p-values at FDR 1% for head and 0.1% for body. Because edges with large p-values are removed, a portion of genes end up being not connected to any other gene in the transcriptome, this resulted in a final dataset of 5261 and 5124 genes for head and body matrices. These FDR thresholds were chosen to reduce the density of the networks as much as possible without removing too many unconnected genes. Since the density of the body graph is higher, we used a more stringent FDR for this tissue. The reduction in density vastly speeds up the computational burden of fitting the subsequent models and allows the analysis to be driven by well-supported edges.

We used the Weighted Nested Degree Corrected Stochastic Block Model (SBM) (Karrer & Newman, 2011; Peixoto, 2017) to cluster genes into blocks. We call gene clusters derived from this approach blocks and not modules because the SBM does not cluster genes following the traditional definition of modularity, i.e., higher within-than between-module correlations, but instead uses a Bayesian approach to maximize the posterior probability of the transcriptome partition given the observed network and edge weights. In this sense, the SBM allows for the discovery of transcriptional structure that is not driven by modularity in its traditional definition. To increase the resolution of the transcriptional partition, we used a nested SBM which clusters genes into blocks in a hierarchical manner, further partitioning blocks into smaller ones for several consecutive levels of the hierarchy (Peixoto, 2017). The nested SBM was fit in graphtools v2.45, using the NestedBlockState model object, with the edge weights given by the arctanh transformed Spearman correlations between gene expressions. The arctanh transformation allows the edge weights to be modeled using normal distributions. The model was fit in three steps: first, an initial partition was obtained with the mcmc_anneal function, which uses Markov Chain Monte Carlo (MCMC, (Peixoto, 2014)) and simulated annealing (Kirkpatrick et al., 1983) to find a partition of the genes into blocks at every level of the SBM hierarchy. Using this initial partition, we then use the mcmc_equilibrate function to find a partition such that subsequent proposals do not improve the posterior probability of the current partition for at least 1000 proposals. The method for fitting the nested SBM using MCMC is described in (Peixoto, 2020). At this point, we consider the block partition is equilibrated and we can use MCMC to sample from the posterior distribution of the block partition. Finally, the posterior sampling is done for 1000 iterations using the mcmc_equilibrate function, and this posterior sample is the partition we use in subsequent analysis. When fitting a nested SBM, the initial annealing step is not always required, but in our case, using this step substantially improved computational performance and allowed us to use the full set of genes in the analysis.

To assess the importance of modularity (as defined above) for transcriptional structure, for each gene block in each level of the hierarchy, we estimated its assortativity, i.e., the contribution of each block to the transcriptome-wide level of modularity (Zhang & Peixoto, 2020) (Figure 4B, Table S3). Assortativity ranges from -1 to 1, with negative numbers indicating that genes in the respective block are less correlated with each other than to genes in other blocks, and therefore that block does not contribute to the modularity of the transcriptome at that level of the hierarchy. Positive numbers indicate that genes in that block behave like modular genes. To see a detailed description of the SBM method, a comparison with traditional modularity-based clustering approaches like WGCNA, and a discussion of its implications for the understanding of transcriptome structure, see Melo et al., 2023)

### Gene connectivity

To determine the level of connectivity of each gene in the transcriptome we used two different metrics estimated using the whole dataset (head samples and genes = 940, 8877; body samples and genes = 939, 8391). First, we estimated the Spearman correlation between all gene pairs, and retained the correlations significant at an FDR 1%. This threshold retained all genes, meaning that all genes retained at least one gene partner after non-significant correlations were removed. Then, we estimated for each gene: a) the number of genes connected to the focal gene (degree), and b) the average correlation between the focal and the connected genes (average correlation).

### GO terms and chromosomal enrichment analysis

ShinyGO v0.77 (Ge et al., 2020) and clusterProfiler v4.2.2 (Wu et al., 2021) were used to calculate the enrichment of gene sets using Biological Process GO terms. The background of genes varies depending on the analysis, e.g., in Figure 1 the background for each tissue corresponds to all genes expressed in that tissue, but for Fig. S1 the background includes only genes expressed in both tissues. The number of enriched GO terms shown and the FDR threshold is indicated in each figure. Chromosomal enrichment was also calculated in ShinyGO using the same background as for GO term enrichment, for this we used sliding windows of 6Mb in steps of 3Mb and the hypergeometric test. Significantly enriched regions are defined as FDR 1e-5.

### Validation of candidate cis-eQTL effects in outbred flies

To validate the effect of *cis-*eQTL on the expression level of the focal gene, we implemented the approach described in (Wolf et al., 2023). We created small outbred populations so that each one was homozygous for one allele at the particular candidate *cis-*eQTL. For this, we randomly sampled hundreds of virgin male and female flies from the same population used for the eQTL mapping (there is a ∼100 generations gap between this and the sampling done for the eQTL mapping experiment). While anesthetized with CO2, we removed one leg per fly and used it for DNA extraction and genotyping. In the meantime, each fly was housed in a separate vial and then mated with individuals of the same genotype.

DNA was extracted following the protocol for QuickEx-tract™ DNA Extraction (cat no. QE09050). The region around the candidate eQTLs was amplified using custom-made primers (Table S8b). Individually barcoded amplicon libraries were prepared using Illumina i5 and i7 primers. Libraries were pooled and sequenced at the Genomics Core Facility of Princeton University using the Miseq nano 150bp PE reads. Genotypes at each focal eQTL were called using bcftools mpileup (Li, 2011) with parameters -Ou -B -q 60 -Q 28 -d 1000 -T -b files | bcftools call -Ou -m -o.

Each population was started from ten individuals of the same focal genotype (five males, five females) which were used to set up five crosses. 10 female and 10 male offspring from each cross were then combined in one bottle where they mated freely for two generations. The resulting populations are the mosaic of ten founder genomes, and share a particular allele at the candidate *cis-*eQTL. Given the large undertaking that this experiment implies, we chose for validation only five ciseQTL-gene pairs (corresponding to three unique SNPs, and to three genes Midway, CG7497, and Ach. Table S8a). Besides being genome-wide significant in the eQTL mapping in both tissues (head and body), the candidate eQTLs have MAF>0.2 to increase the probability that a random sample of the population will contain some flies homozygous for the minor allele, are associated with only one gene in cis (but could be regulating different genes in each tissue), and have large DeepArk predicted effects >10%.

### RNA extraction and qPCR

For each population, four pools of seven-day old female flies (mated) were collected. If the candidate eQTL had an effect in head tissue, each pool consisted of 11 heads, if the effect was to be tested in the body, each pool consisted of three bodies. Gene expression was only quantified in females to match the eQTL mapping experiment. Each pool of heads or bodies was placed in a well of 96-well plates and total RNA was extracted using Quick-RNA 96 Kit by Zymo Research (Catalog no. R1053). 350 ng of RNA were used as input for cDNA synthesis using SuperScript™ III First-Strand Synthesis System by Invitrogen (Catalog no. 18080051), and rt-qPCR (using PowerTrack™ SYBR Green Master Mix by Applied Biosystems Catalog no. A46012) was used to quantify gene expression in each pool. The rt-qPCR primers used to amplify the target genes as well as the reference gene rpl32 are shown in Table S8c. The rt-qPCR assay was run in a 384-well plate that included all samples for all candidate eQTL-gene pairs.

#### Data analysis

Raw qPCR data without baseline correction was imported into LinRegPCR v2020.0 (Ramakers et al., 2003; Ruijter et al., 2009) where the PCR efficiency for each primer pair was estimated. Deviation from the expected PCR efficiency of two is known to cause large biases in the estimation of fold changes in gene expression (Ramakers et al., 2003; Ruijter et al., 2009). LinRegPCR uses linear regression of log (fluorescence) to estimate the PCR efficiency per sample, and uses the average efficiency of all samples per amplicon group (i.e., per primer pair) to estimate the amount of starting RNA concentration per sample (N0) in arbitrary fluorescence units. The amount of starting RNA per sample is then estimated as the ratio of N0 for the target gene (gene affected by eQTL) and N0 for the reference gene (rpl32). The difference in gene expression levels between genotypes at a particular eQTL was tested using t.test (see Table S8a for results and sample size).

