## Supplementary methods and figures for "Saturating the eQTL map in *Drosophila melanogaster*: genome-wide patterns of cis and trans regulation of transcriptional variation in outbred populations"

**This PDF file includes:**

Materials and Methods

Figures S1 to S19

Tables S1 to S8 are provided in a separate spreadsheet

### Materials and Methods

#### mRNA extraction and RNAseq library preparation

We used a previously optimized low-cost and high-throughput protocol for mRNA extraction from single fly heads (Pallares et al., 2020); the protocol is described in detail here: Suppl. File 2 of (Pallares et al., 2020). Briefly, head and body samples were processed in the original 96-well plates in which they were previously frozen. The tissue was mechanically homogenized in a Talboys High Throughput Homogenizer (#930145) using one 2.8 mm stainless-steel bead (OPS diagnostics, #089- 5000-11) and 100 ul of lysis buffer per well. The lysate was transferred to a new 96-well plate and mRNA extraction was performed using the CyBio® FeliX pipetting robot (Analitik Jena) using the Dynabeads™ mRNA DIRECT™ Purification Kit (ThermoFisher, #61012). Dynabeads are oligo-(dT) beads that bind the poly-adenylated tails of RNA molecules. For single head mRNA extraction, the final elution was done in 10ul 10mM TRIS-HCl resulting in ~10-20 ng of mRNA per head. For single body extraction, the final elution volume was 30 ul resulting in 90-180 ng of mRNA per body.

RNAseq library preparation was done following the TM3'seq protocol (Pallares et al., 2020); a detailed description of the protocol, primers, and reagents is available at https://lufpa.github.io/TM3Seq-Pipeline/tm3seq_protocol and in Suppl. File 1 of Pallares et al. Briefly, during first strand synthesis, adapter B was ligated to the polyA tail of the RNA molecule; this adapter will be used to ligate the sequencing adapter and i7 barcode during the final amplification step. Using home-brew Tn5 transposase, the cDNA was tagmented at the same time the adapter A was ligated to the cDNA fragments. In the final amplification step, Illumina i7 and i5 primers were used to amplify cDNA fragments that contained both, adapter B and adapter A, assuring that only the 3' end of the molecule is amplified.

Individual libraries processed in the same plate were pooled (2ul per body library, 5ul per head library) and each pool was cleaned and size-selected for a mean size of ~300-400bp using double-sided cleanup approach (0.6x right - 1x left) with Agencourt AMPure XP beads (Beckman Coulter). The quality and concentration of each pool was assessed using Pooled Agilent TapeStation before combining pools for sequencing. Samples were sequenced using 150bp SE reads on the Illumina NovaSeq S4 platform at the New York Genome Center and Illumina NovaSeq S2 at the Genomics Core Facility of the Lewis-Sigler Institute for Integrative Genomics at Princeton University.

Protocols and subroutines for mRNA extraction, cDNA synthesis, and library preparation were implemented in the CyBio® FeliX liquid handling robot and are available upon request.

#### DNA extraction and library preparation

DNA was extracted from the body or head lysate used to first extract mRNA in the following way: after mRNA bound to the Dynabeads™ in step 13 of the RNA extraction protocol (see https://lufpa.github.io/TM3Seq-Pipeline/mrna_extraction), the supernatant was transferred to a deep 96-well plate and mixed with 400ul of genomic lysis buffer (Zymo, #D3004-1). The mix was stored at -20C to await further processing. Samples were transferred to an Acroprep advance 1mL DNA binding plates (Pall Life Sciences, #8132) for DNA extraction using the Multi-Well Plate Vacuum Manifold (Pall Life Sciences, #5017) and DNA pre-wash, and gDNA wash buffers from Zymo (#D3004-1, #D3004-5). DNA library preparation was also performed in 96-well plates, following the tagmentation approach outlined in Picelli et al. (2014) and implemented in a CyBio® FeliX liquid handling robot (Analitik Jena). In a first step, 10ul (100uM) of forward oligo adapter A (or adapter B) and 10ul (100uM) of reverse oligo Tn5MErev were mixed with 80µl reassociation buffer (10mM Tris pH 8.0, 50mM NaCl, 1mM EDTA) and annealed in thermocycler using the following program: 95 °C 10 min, 90 °C 1 min, reduce temperature by 1 °C/cycle for 60 cycles, hold at 4 °C. Second, the pre-annealed adapters A and B were loaded onto the home-brewed Tn5 transposase by mixing 45ul of Tn5 with 9ul each of pre-annealed adapter A and B (10uM) and incubating the mix in a thermocycler for 30 min at 37 °C. Finally, the charged Tn5 was diluted 1:1 with reassociation buffer:glycerol (1:1). The DNA tagmentation was performed by incubating for 7 min at 55 °C a mix of 5ul DNA sample, 1ul of diluted Tn5, 2µl of 5X TAPS buffer pH 8.5 (50mM TAPS, 25mM MgCl2, 50% v/v DMF), and 5µl of water. Then, 2.5µl of 0.2% SDS (Promega, #V6551) was added to the mix and incubated in a thermocycler for 7 min at 55 °C to dissociate the Tn5 from the DNA. The final step of PCR library amplification was performed by combining 2µl of the tagmentation reaction with 7µl of OneTaq HS Quick-Load 2x (NEB, #M0486L), 1µl of the i5 primer (5uM), 1µl of the i7 primer (5µM), and 4µl of water. The mix was amplified on a thermocycler using the following program: 68 °C 3 min, 95 °C 30sec, [95 °C 10sec, 55 °C 30sec, 68 °C 30sec] for 18 cycles, 68 °C 5 min.

For each plate, 2µl per well of the final PCR amplification reactions were pooled, cleaned, and size selected for an average size of 400bp using a double-sided cleanup approach (0.6x-1x) with Agencourt AMPure XP beads (Beckman Coulter). Library size and concentration were checked for each pool using Agilent TapeStation before pooling multiple plates for sequencing. Samples were sequenced using 100bp PE reads on the Illumina NovaSeq S4 platform at the New York Genome Center.

### eQTL mapping

To identify genetic variation regulating transcript abundance, we used GEMMA (X. Zhou & Stephens, 2012). In contrast with Matrix eQTL (Shabalin, 2012) which has been explicitly designed for eQTL mapping, GEMMA is much slower and requires post-hoc processing of cis-eQTL and trans-eQTL results. However, given the nature of our experimental design, where each female fly can lay many eggs during each round of egg lay, we expect a wide range of relatedness between individuals (see Fig. S2). GEMMA provides the flexibility to fit both fixed and random effect models (Mixed Linear Models - MLM) that account for variation in relatedness between individuals using a genetic relationship matrix (GRM). Matrix eQTL, on the other hand, implements a linear model, the GRM effect is the same for all genes resulting in an overly conservative correction of the effect of relatedness between individuals. Note that the GRM effect is the same for all genes, resulting in an overly conservative correction of the effect of relatedness between individuals. In Fig. S19 we show the comparison between GEMMA and Matrix-eQTL results for a subset of 199 randomly selected genes measured in 856 fly bodies. GEMMA recovers 99.25% of cis-eQTLs and 86.5% of trans-eQTLs detected with Matrix-eQTL while finding many more eQTLs that have been penalized in Matrix-eQTL. For contrast, we also show the results from Matrix-eQTL when relatedness between samples is not incorporated in the model.

### Supplementary Figures

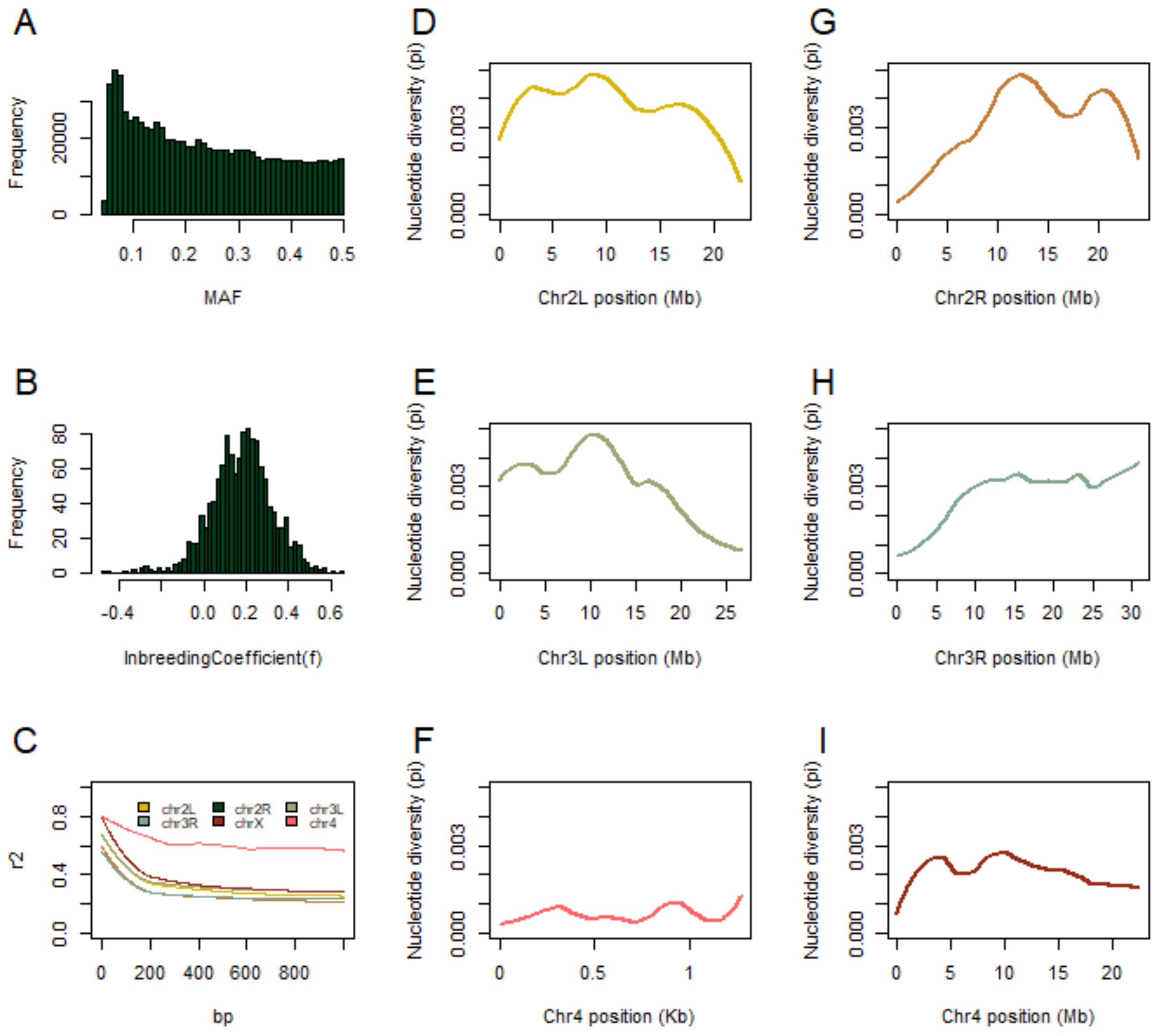


**Figure S1. Population-level parameters for the Nex fly population. (A-C)** Minimum allele frequency distribution (MAF), inbreeding coefficient (f), and linkage disequilibrium ($r^2$) were estimated on the full dataset (n=1286 fly samples) and the full set of 854,350 SNPs with MAF>0.05 before LD pruning. **(D-I)** Nucleotide diversity (pi) was estimated using a sample of the 320 best-covered samples (coverage >8x) in 1Kb windows. Each chromosome is shown in a separate plot and the lines correspond to Loess regressions with smoothing parameter alpha = 0.5. Average pi per chr: 2L=0.0038, 2R=0.0032, 3L=0.0031, 3R=0.0027, 4=0.00058, X=0.0021.

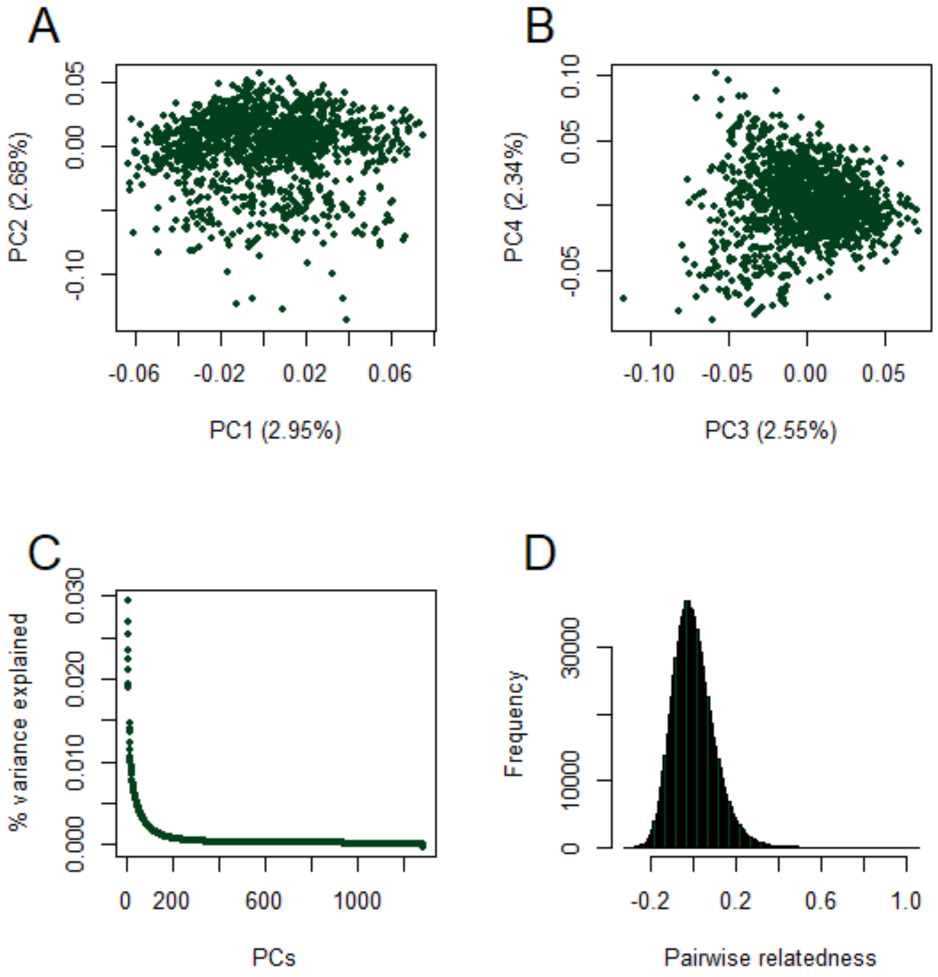


**Figure S2. Genetic structure in the Nex mapping population.** 1286 samples and 389,896 LD-pruned SNPs were used to estimate the genetic relationship matrix (GRM). **(A, B)** The first four PCs from the GRM and **(C)** the amount of genetic variation explained by each PC are shown. **(D)** Pairwise relatedness between all samples was estimated using the full set of quality-filtered SNPs before LD pruning (n=854,350).

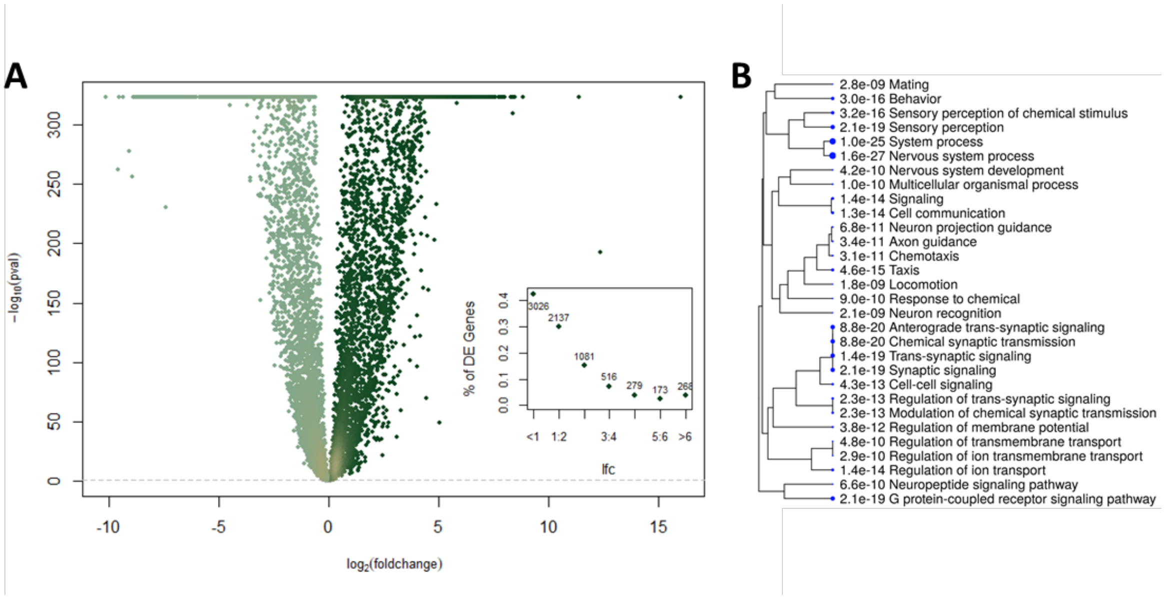


**Figure S3. Patterns of differential gene expression.** 7795 genes are expressed in both tissues, head and body, and 94% of those are significantly differentially expressed (n=7374). **(A)** Volcano plot showing the fold change (lfc) and significance of the difference in mean gene expression between head and body samples. Each dot represents a gene, dark green indicates genes with higher mean expression in the head, and light green genes with higher expression in the body. High density regions are highlighted in yellow. Genes above the dotted line are significantly differentially expressed at FDR 5%. The inset shows the number of genes for several lfc categories. **(B)** GO Biological Process enrichment analysis of the top 1000 differentially expressed genes ranked by lfc. The tree shows the top 30 GO terms clustered based on the number of genes shared, and the p-value of the enrichment. Analysis done in ShinyGO v0.741 (Ge et al., 2020).

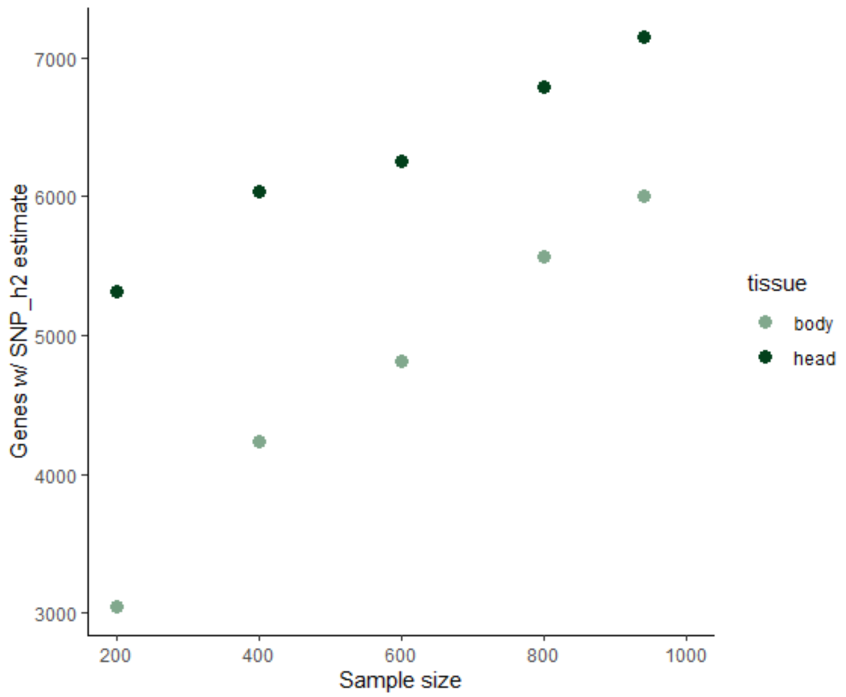


**Figure S4. Effect of sample size on the number of genes for which SNP heritability can be estimated.** The full dataset for each tissue was randomly down-sampled to sets of 200 to 800 samples, and SNP_h2 was estimated for each gene in the dataset using GEMMA (Zhou & Stephens, 2012). We define non-estimable SNP_h2 as values that hit the lowest boundary of GEMMA estimates of 9.99999e-6 (see Methods section 'Heritability estimates'). For each subset of samples, only genes passing the expression filters were used (i.e., average CPM>1 and detected in at least 20% of the samples). The number of genes per dataset are shown in Table S1, it ranges from 8331 genes in the dataset of 200 body samples to 8877 genes in the full dataset of head transcriptomes.

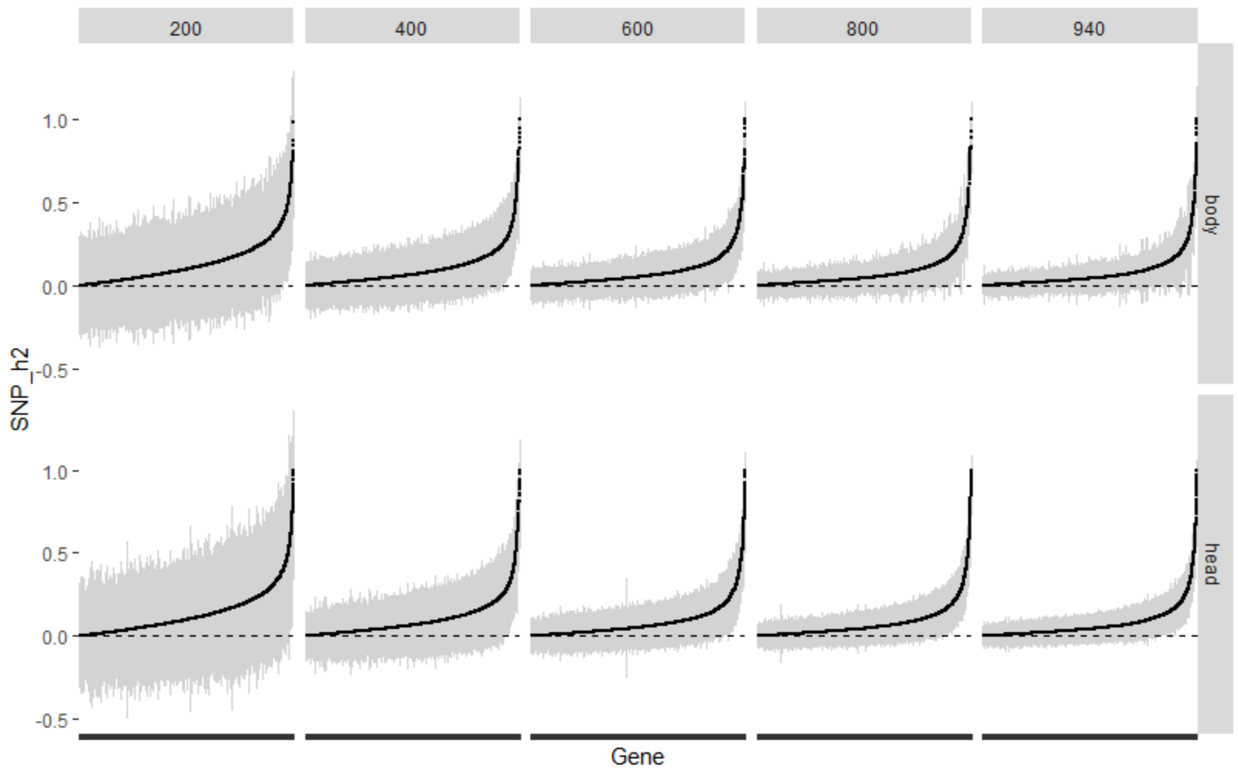


**Figure S5. SNP heritability estimates for different sample sizes.** The full dataset for each tissue was randomly down-sampled to sets of 200 to 800 samples, and SNP_h2 was estimated for each gene in the dataset using GEMMA (Zhou & Stephens, 2012). Point estimates (dots) and two standard errors around the estimate (grey lines) are shown for each dataset in each tissue. For each subset of samples, only genes passing the expression filters were used (i.e., average CPM>1 and detected in at least 20% of the samples). The number of genes per dataset are shown in Table S1, it ranges from 8331 genes in the dataset of 200 body samples to 8877 genes in the full dataset of head transcriptomes.

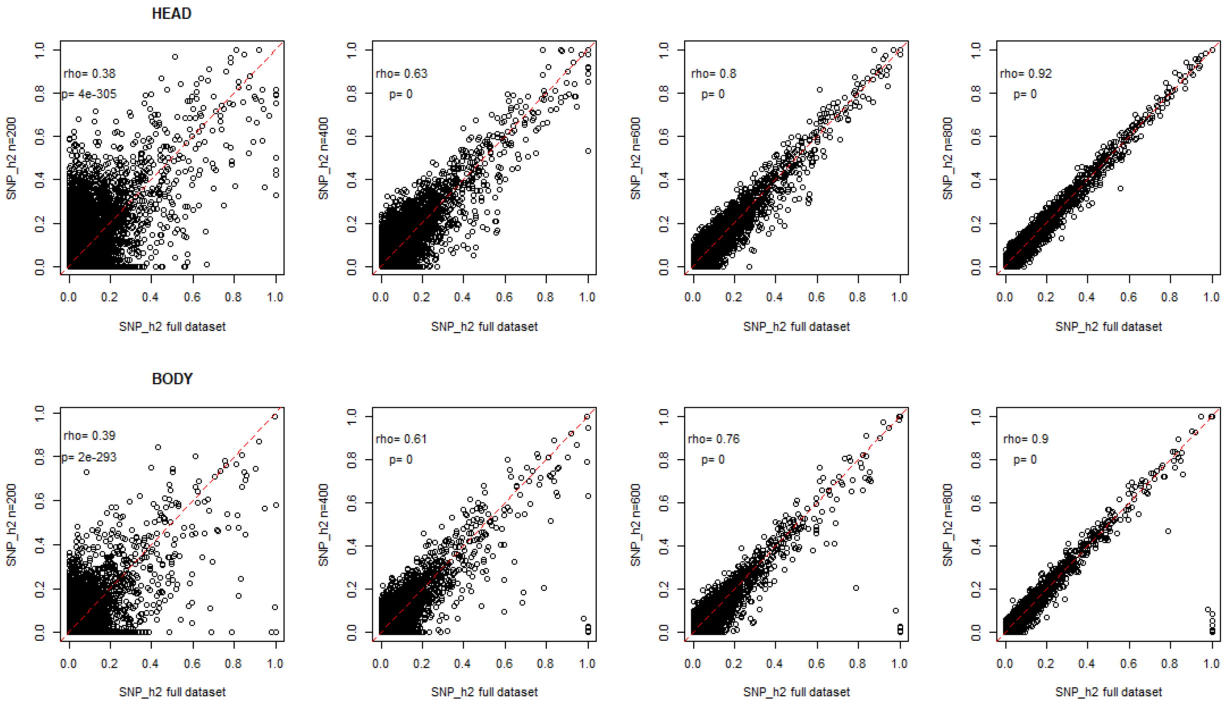


**Figure S6. Correlation between SNP heritability estimates using the full dataset (x axis) and smaller subsets of data (y axis)**. The upper row shows heritability estimates done on head-derived transcriptomes; the lower row corresponds to body transcriptomes. Spearman correlation (rho) and p-value are shown for each comparison.

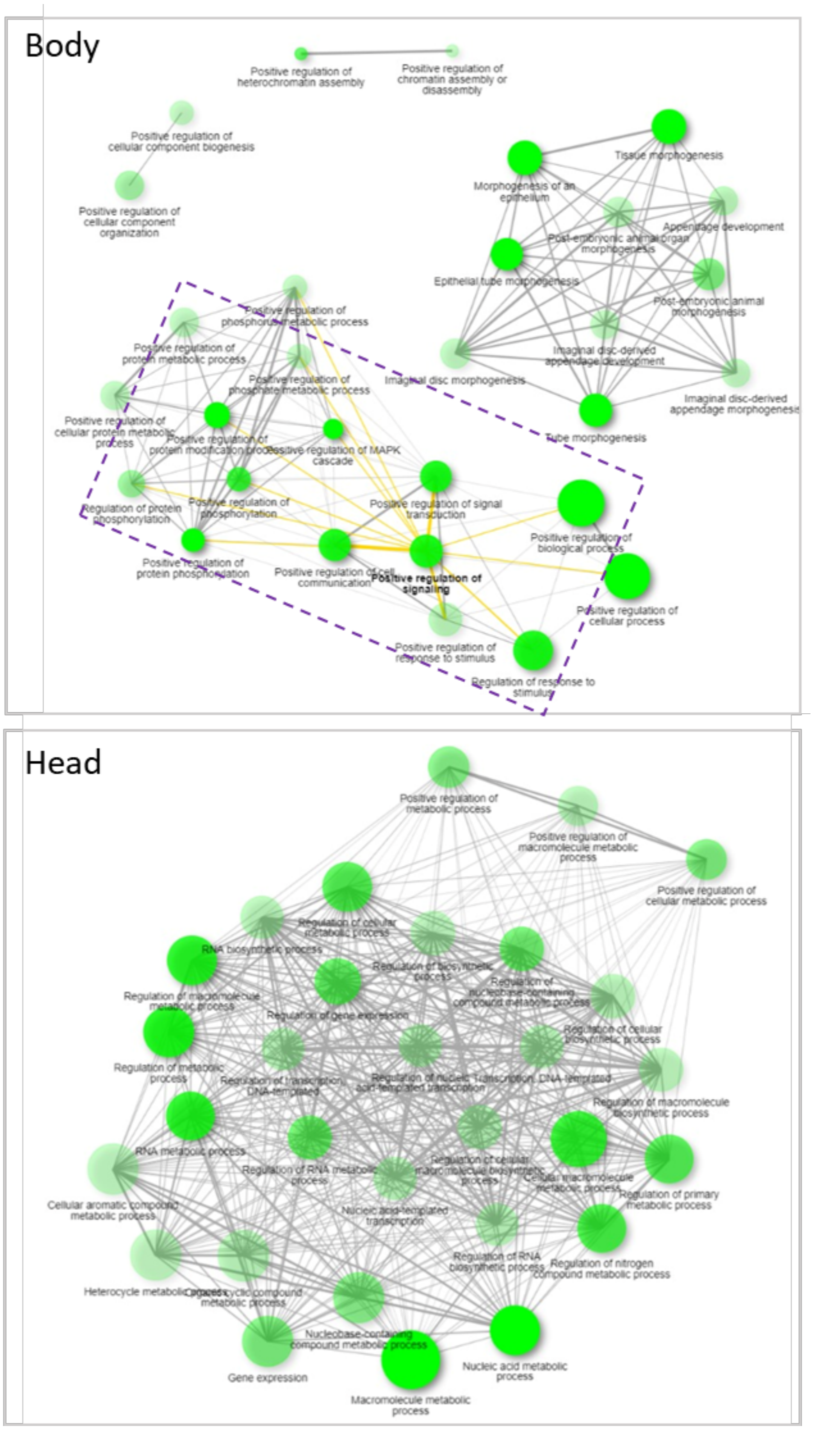


**Figure S7 GO enrichment (Biological Process) of low heritability genes.** Genes were divided in five quantiles based on their SNP_h2; GO enrichment was estimated for genes in the first quantile in body (n = 1201) and head (n = 1431) using ShinyGO v0.741 (Ge et al., 2020). Each dot is an enriched GO term, dot size corresponds to the number of genes in the GO term, and darker colors indicate lowest enrichment p-values. Edges represent the number of genes shared between dots and are drawn when there are 20% or more shared genes. The top 30 enriched GO terms at FDR 1% are shown. The dashed rectangle indicates GO terms in body also found in the highly connected head network.

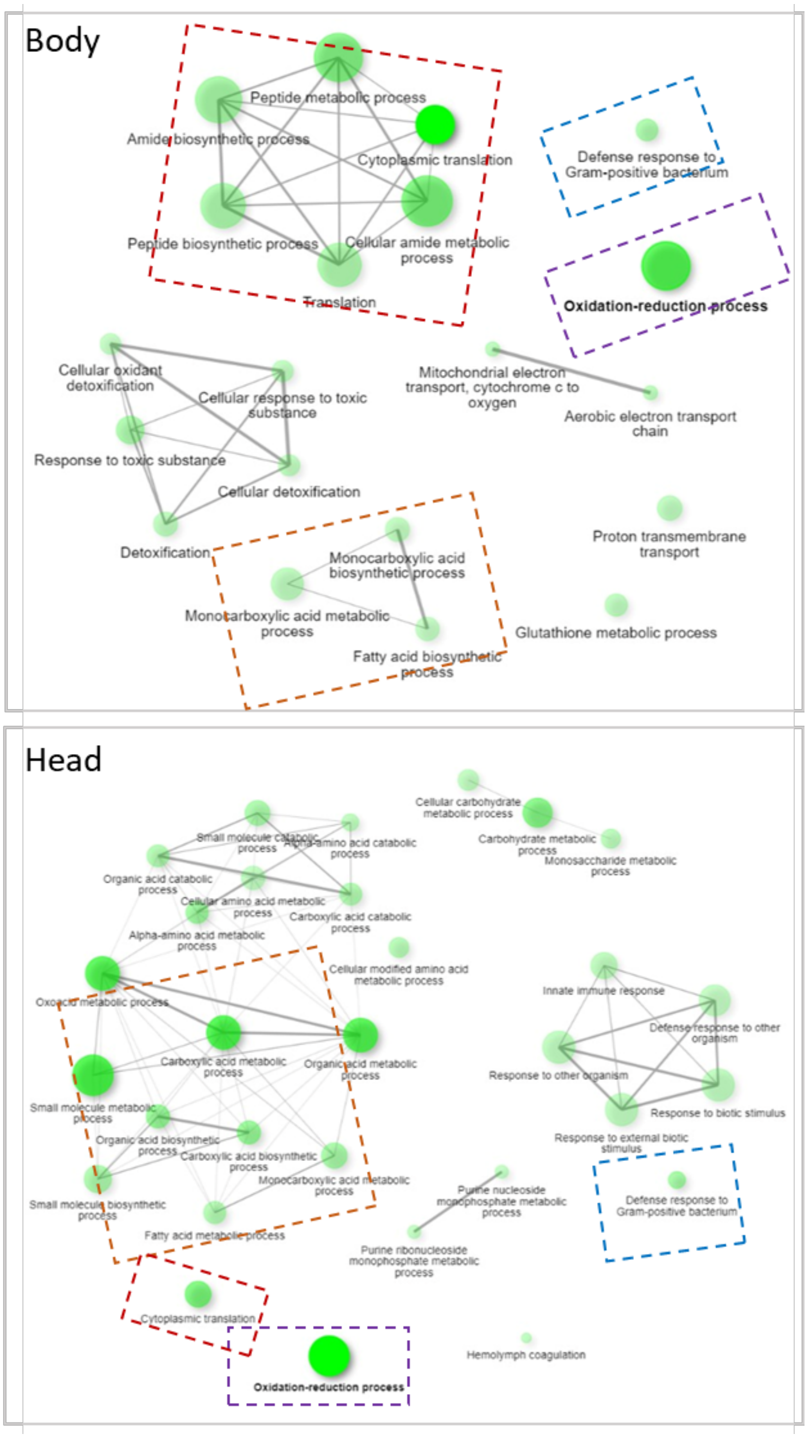


**Figure S8. GO enrichment (Biological Process) of high heritability genes.** Genes were divided in five quantiles based on their SNP_h2; GO enrichment was estimated for genes in the fifth quantile in body (n = 1200) and head (n = 1431) using ShinyGO v0.741 (Ge et al., 2020). Each dot is an enriched GO term, dot size corresponds to the number of genes in the GO term, and darker colors indicate lowest enrichment p-values. Edges represent the number of genes shared between dots and are drawn when there are 20% or more shared genes. The top 30 enriched GO terms at FDR 1% are shown. Dashed rectangles with matching colors highlight modules that are present in both tissues.

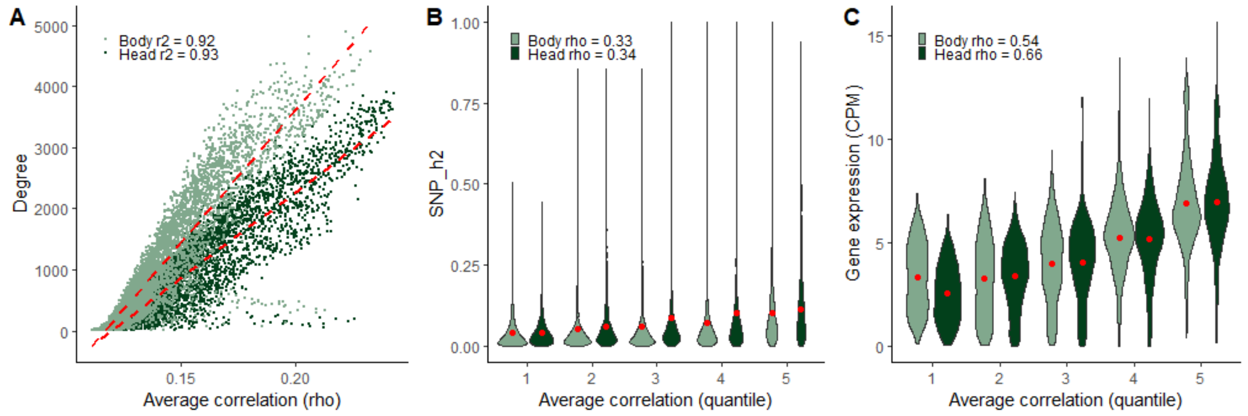


**Figure S9. Gene-level connectivity measures. (A)** Rank correlation between degree (number of genes the focal gene is connected to) and average correlation with all other genes in the transcriptome. Relationship between average correlation and **(B)** SNP_h2 and **(C)** mean gene expression (log2 CPM). Correlations are shown for each comparison, p-value in both cases < 2.2e-16. Average Spearman correlation (rho) and degree are estimated only for gene pairs whose correlation is supported at FDR 1%. Plots similar to **B** and **C** but using degree instead of average correlation as the connectivity variable are shown in Fig. 3. Only genes with non-zero heritability estimates were used here n(head)=7155, n(body)=6001.

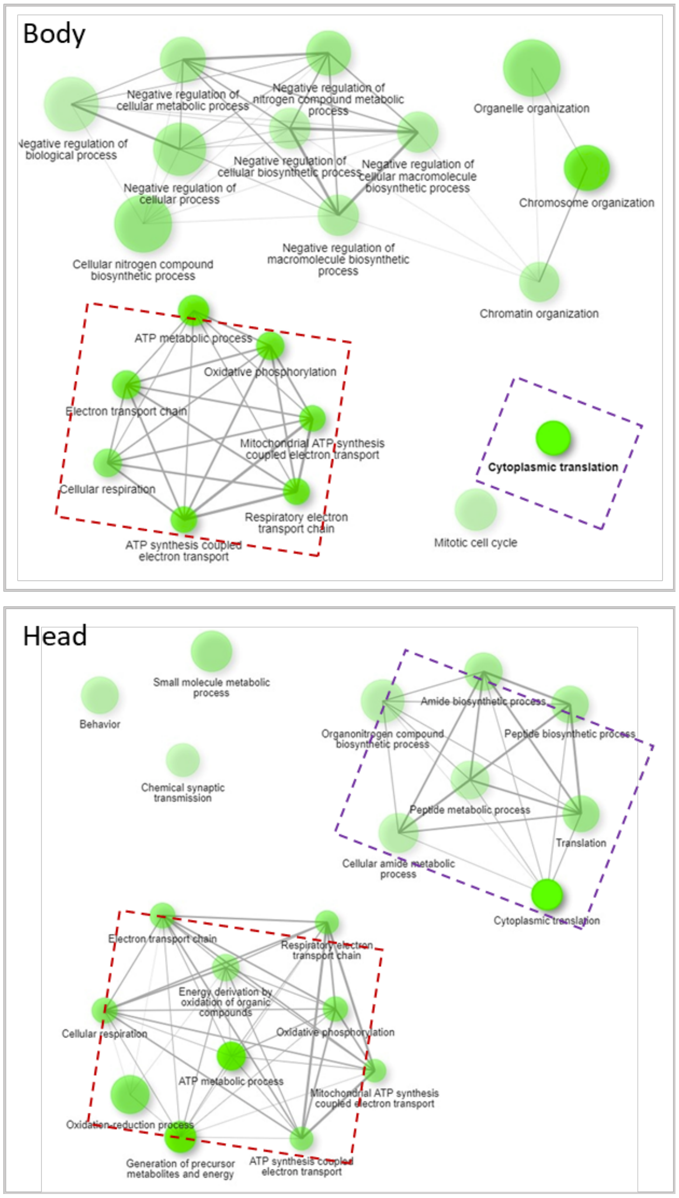


**Figure S10. GO enrichment (Biological Process) of highly connected genes.** Genes were divided in five quantiles based on their degree (i.e. number of genes the focal gene is connected to, based on Spearman correlation p-values at FDR1%); GO enrichment was estimated for highly connected genes (i.e. genes in the fifth quantile, median degree: 1201 body, 1431 head) using ShinyGO v0.741 (Ge et al., 2020). Each dot is an enriched GO term, dot size corresponds to the number of genes in the GO term, and darker colors indicate lowest enrichment p-values. Edges represent the number of genes shared between dots and are drawn when there are 20% or more shared genes. The top 30 enriched GO terms at FDR 1% are shown. The dashed rectangle indicates GO terms shared between head and body.

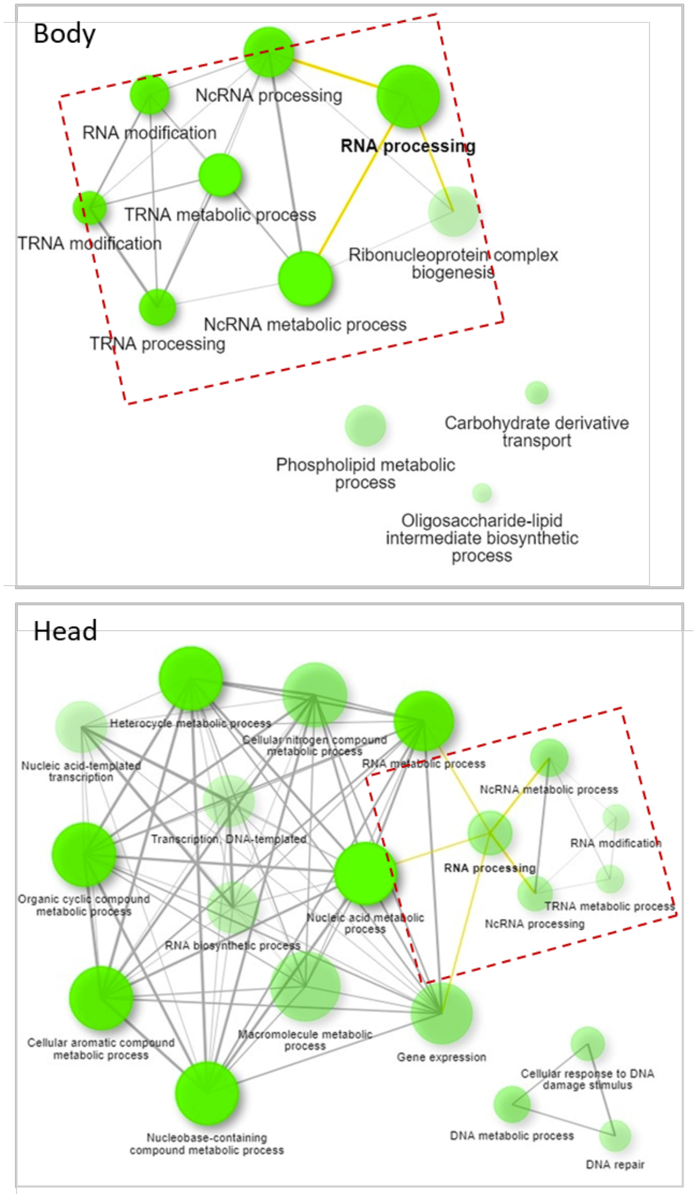


**Figure S11. GO enrichment (Biological Process) of poorly connected genes.** Genes were divided in five quantiles based on their degree (i.e. number of genes the focal gene is connected to, based on Spearman correlation p-values at FDR1%); GO enrichment was estimated for lowly connected genes (i.e. genes in the first quantile, median degree: 60 body, 37 head) using ShinyGO v0.741 (Ge et al., 2020). Each dot is an enriched GO term, dot size corresponds to the number of genes in the GO term, and darker colors indicate lowest enrichment p-values. Edges represent the number of genes shared between dots and are drawn when there are 20% or more shared genes. The top 30 enriched GO terms at FDR 1% are shown. The dashed rectangle indicates GO terms shared between head and body.

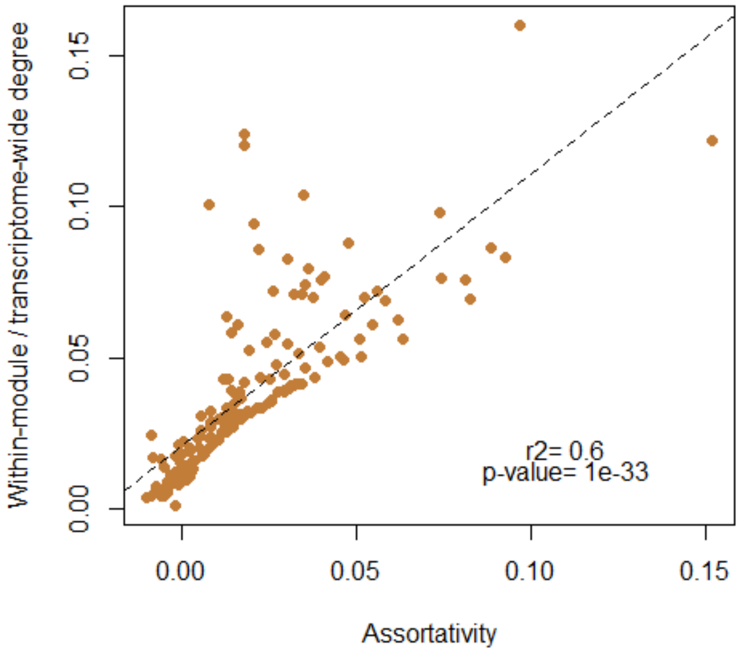


**Figure S12. Correlation between connectivity and assortativity**. For each gene block the average within-block degree was estimated (average number of genes a focal gene is connected to inside the module of the focal gene. Only Spearman correlations at FDR 10% were included), as well as the total average degree (average number of genes the focal gene is connected to in the whole transcriptome. Only Spearman correlations at FDR 10% were included). The ratio between within-module and transcriptome-wide connectivity measures was estimated as a proxy for the traditional definition of modularity which implies stronger correlation within that between modules (degree and Spearman correlation are highly correlated, see Fig. S7). The high correlation between the ratio of within-module/total connectivity and assortativity ($r = 0.776$, p-value $< 2e-16$,) shows that assortativity recapitulates the traditional meaning of modularity.

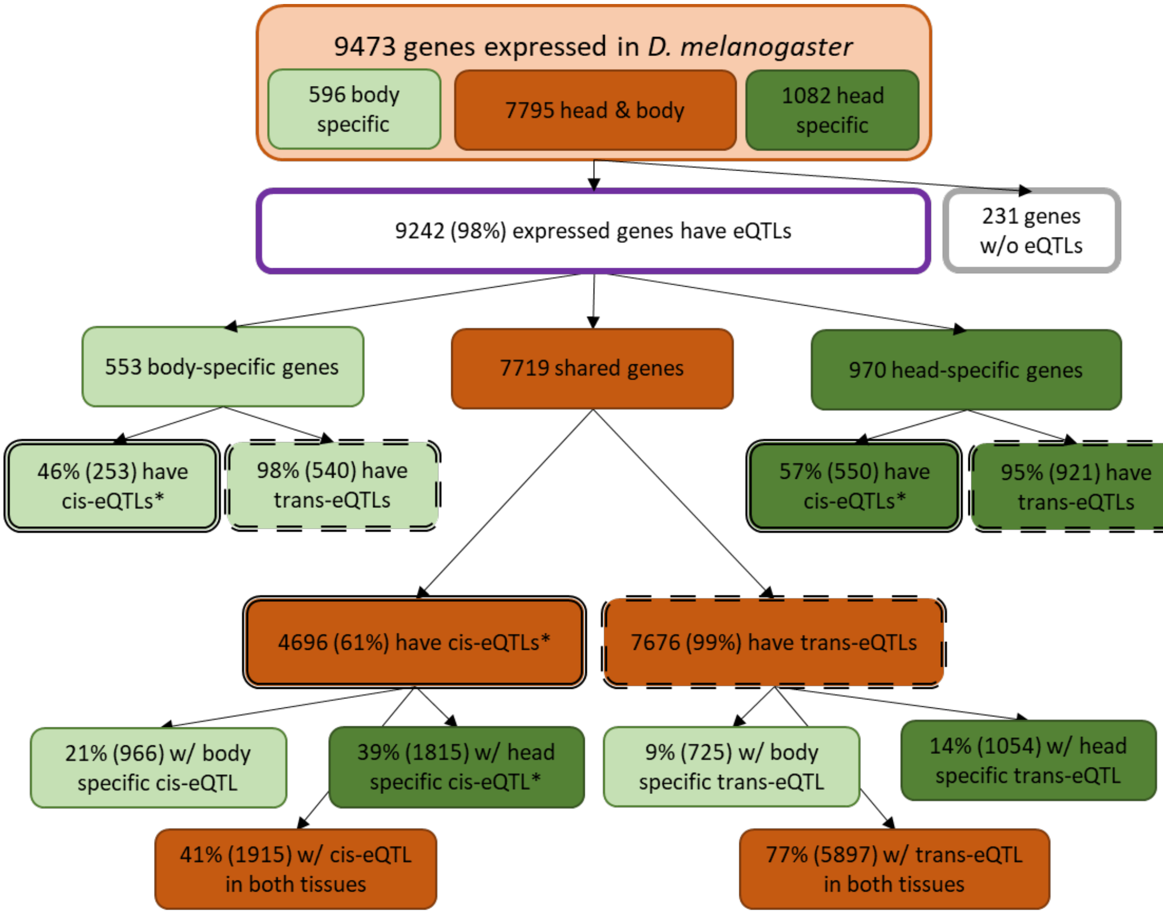


**Figure S13. Gene expression regulation in outbred *D. melanogaster***. 9473 genes are expressed in female fruit flies. A gene is considered expressed in a tissue (head or body) if it is present in at least 20% of the individuals and has an average CPM>1. eQTLs were identified for 98% of the genes expressed in the fruit fly (n = 9242), of those, 60% have at least one cis-eQTL (5499 genes, double line) and 99% have at least one trans-eQTL (9137 genes, dashed line). The diagram shows the expression pattern of each gene (orange: shared, or tissue-specific) and the proportion of such genes with cis and trans-eQTLs. *The number of genes with cis-eQTLs could be a slight underestimate because not all genes had cis-SNPs in the panel of SNPs used for mapping. However, for the sake of simplicity in this figure the percentages are calculated using the total number of genes (n genes tested = 8877 head, 8391 body; n genes with cis-SNPs = 8834 head, 8357 body). Most genes expressed in both tissues (77%) have trans-regulation in both tissues, while only 40% have cis-regulation in head as well as body.

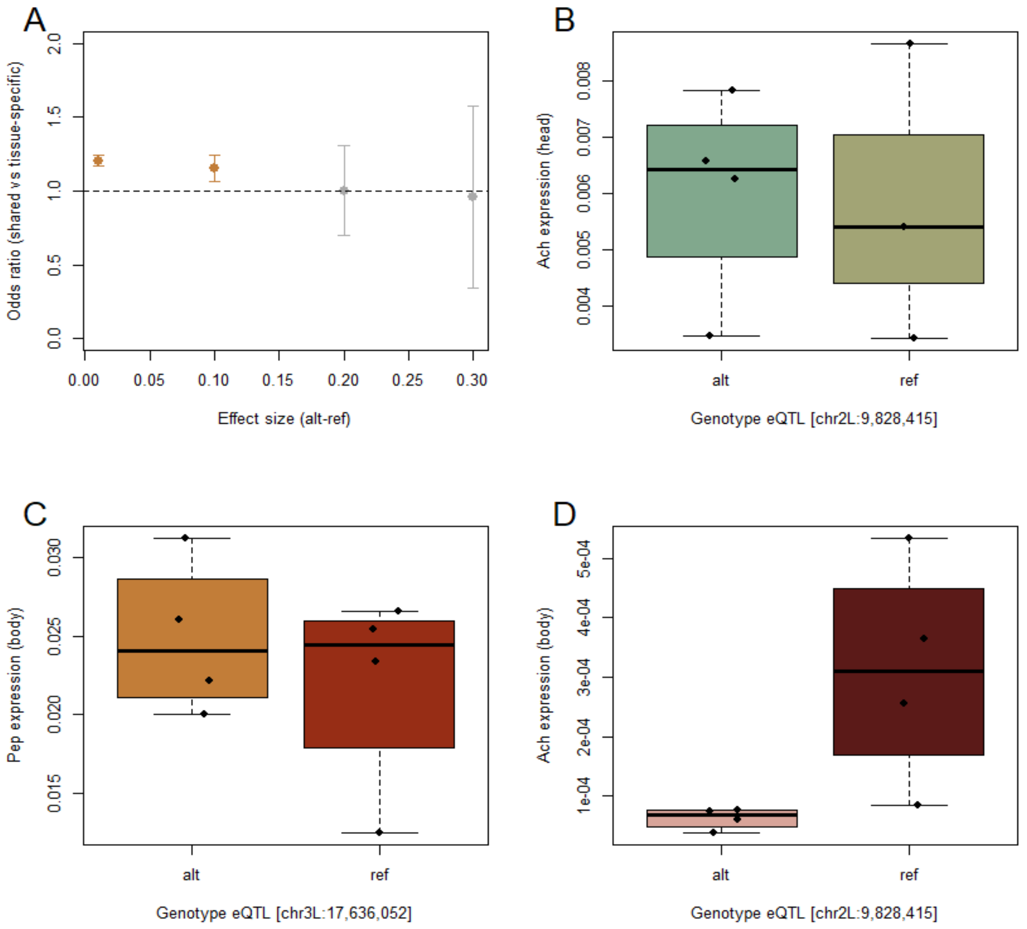


**Figure S14. Validation of candidate eQTL. (A)** cis-eQTL that were identified in both tissues, head and body, are enriched for large predicted effects compared to tissue-specific cis-eQTL. The predicted effect was estimated with the deep-learning approach implemented in DeepArk (Cofer et al., 2021). Sample size of comparisons: shared eQTL = 15,472; tissue-specific eQTL = 53,387. For each effect size, groups were compared using Fisher exact tests. All significant tests (orange) have p-values < 0.001. Grey comparisons were not significant at p-value 0.05. 95% confidence intervals for odds ratio are shown. **(B-D)** Experimental validation of cis_eQTL effects. Five candidate eQTL-gene pairs were experimentally validated using rt-qPCR (see Methods section "Validation of candidate cis-eQTL effects"), the results for the genes *Midway* and *CG7497* are shown in Figure 7. Here we show the gene expression effect of three additional eQTL-gene pairs. The genotype at the candidate eQTL doesn't seem to affect the expression levels of the *Ach* gene in head tissue (**B,** t.test p-value = 0.91, fold change 1.04x) and the *Pep* gene in body tissue (**C**, t.test p-value = 0.51) while it seems to regulate expression of the *Ach* gene when expressed in the body (**D**, t.test p-value 0.078). When the eQTL effect was detected in head tissue in the initial eQTL mapping, gene expression was quantified in four pools of 10 heads each. When the body was the focal tissue, 4 pools of 3 bodies were used for expression quantification.

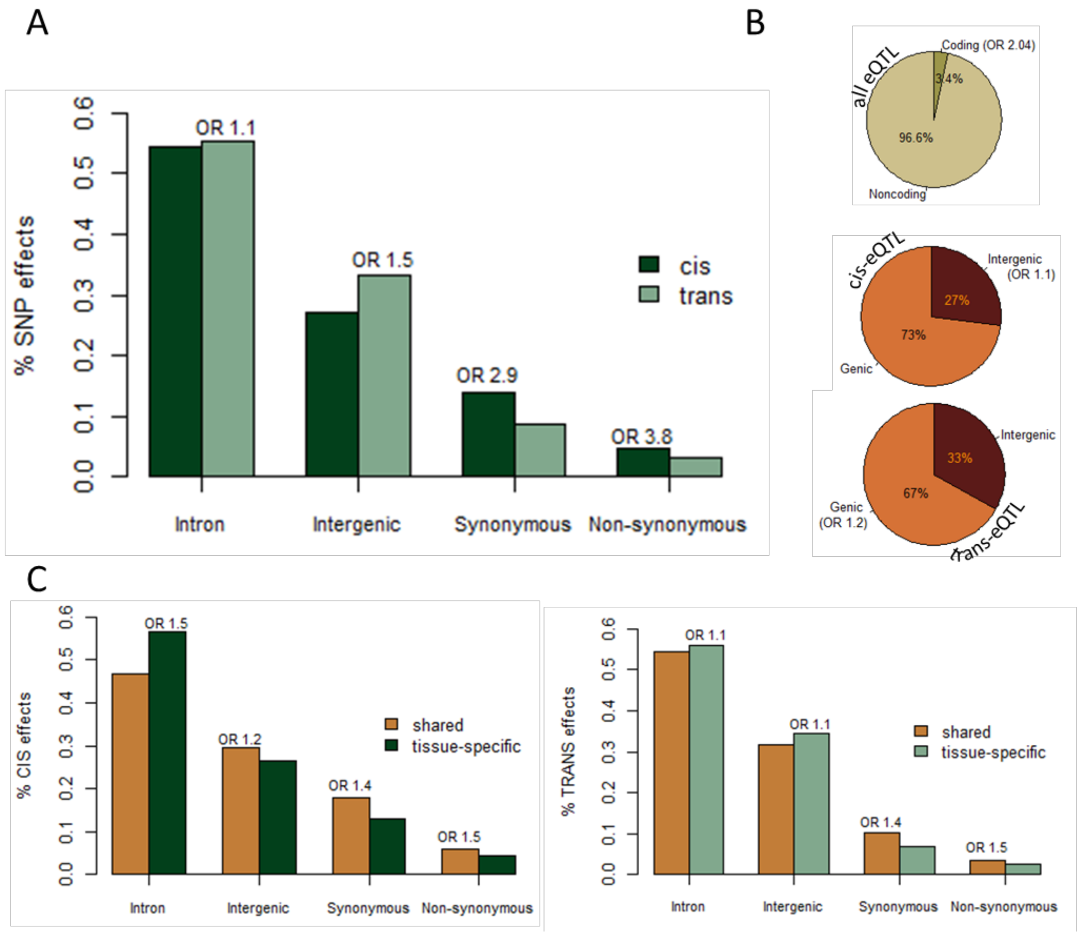


**Figure S15. eQTL annotation.** The genomic location and predicted effect on protein sequence is shown for cis and trans-eQTLs **(A)**, as well as for shared and tissue-specific eQTLs **(C)**. When there is significant enrichment for a certain category, the odds ratio is shown on top of the bar corresponding to the enriched group. **(B)** Comparison between eQTLs and genomic background. Cis and trans-eQTLs are enriched for coding effects compared to non-significant SNPs. Because cis and trans values are very similar (Non-coding eQTLs: cis 95%, trans 97%) an average of the two was used for visualization. Odds ratio also corresponds to the average between cis and trans (chi-square cis p<2e-16, odds ratio 2.0; trans p<2e-16, odds ratio 1.9). Cis eQTLs are enriched in intergenic effects relative to non-cis SNPs (cis p <2e-16, odds ratio 1.1) while trans-eQTL are enriched in genic location relative to non-trans SNPs (trans p <2e-16, odds ratio 1.2).

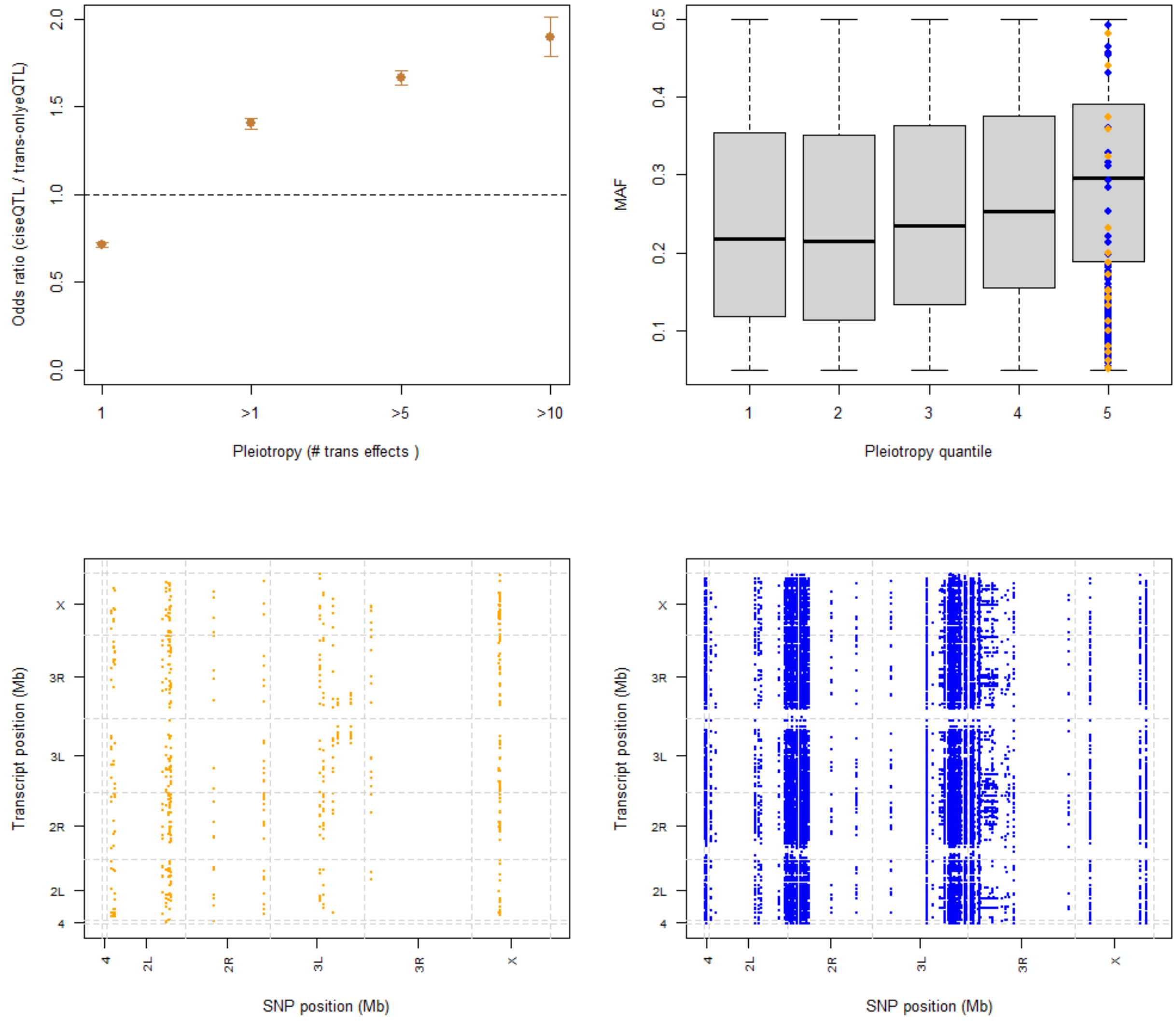


**Figure S16. Trans-regulation hotspots.** Most eQTL affect one or very few genes, however there are some, called here hot-eQTL (affecting more than 20 genes), that have tens or hundreds of trans effects. **(A)** cis-eQTL are significantly more likely to be pleiotropic (number of trans-effeccts >1) than trans-only-eQTL. While eQTL that act only in trans are significantly more likely to affect just one gene (number of trans-effects =1). Information about the Fisher tests, p-values and number of eQTL per group are found in Table S8. All Fisher tests are significant at p-value 0.05. **(B)** The minor allele frequency distribution (MAF) of eQTL with different levels of pleiotropy is not significantly different. Hot-eQTL are defined here as eQTL in the fifth quantile of pleiotropy span the entire allele frequency distribution (orange = hot-eQTL in head, blue= hot-eQTL in body). **(C, D)** Dotplot highlighting hot-eQTLs that affect more than 20 genes in trans (Orange, 22 hot-eQTL in head; blue, 189 in body) and their distribution across the genome.

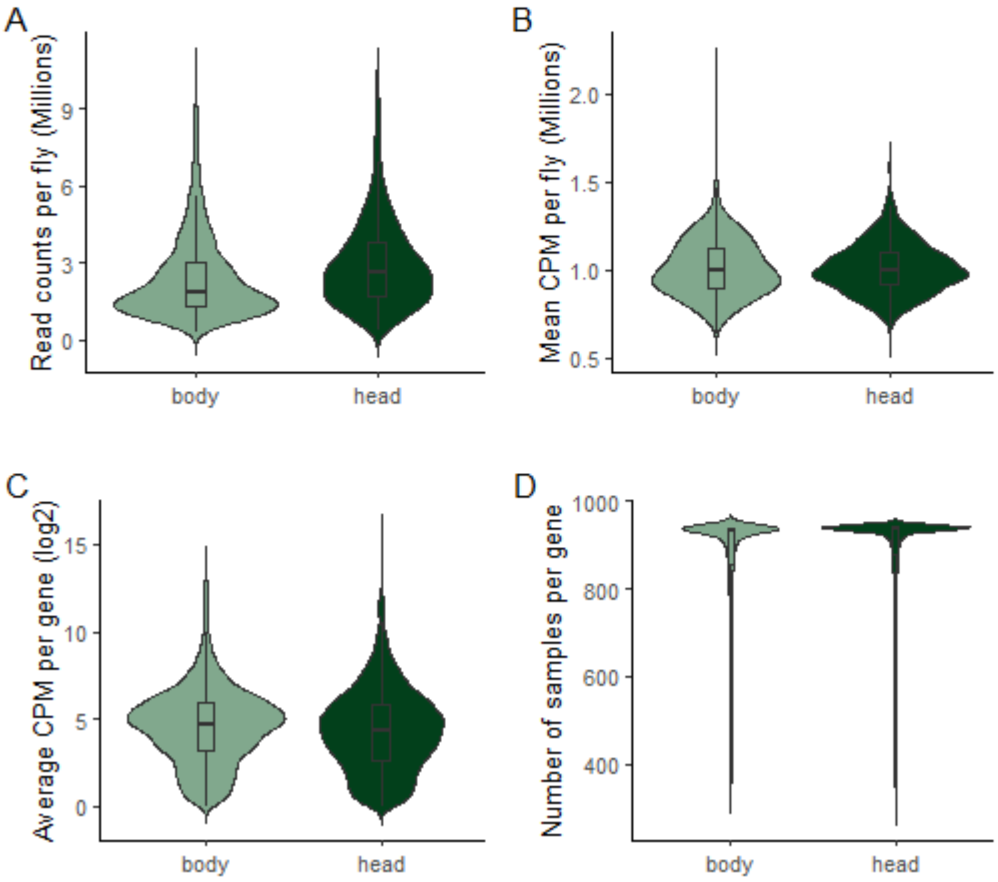


**Figure S17.** RNAseq samples. Coverage parameters per fly **(A, B)** and gene **(C, D)** are shown. Coverage per fly in **(A)** raw and **(B)** normalized counts is shown, as well as **(C)** average normalized counts per gene and **(D)** number of fly samples in which a gene was detected (raw counts > 0).

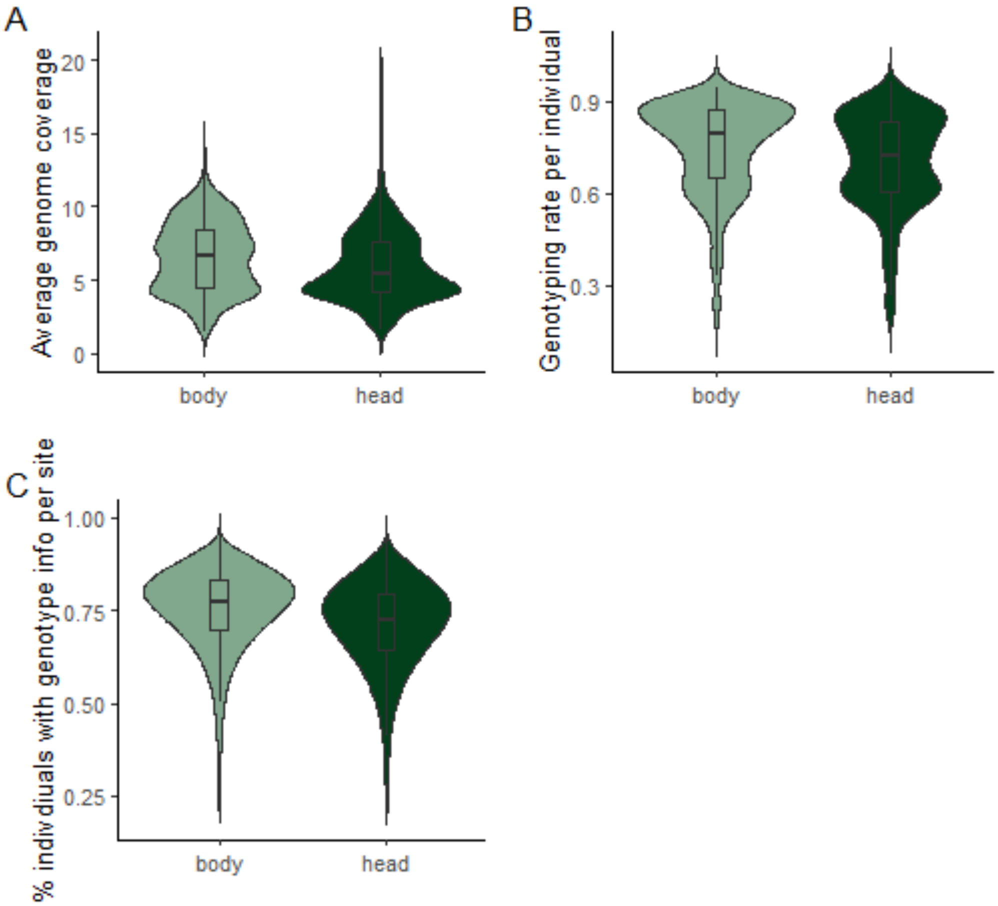


**Figure S18.** DNAseq samples. **(A)** Genome coverage, **(B)** percentage of total SNPs (n=389,896) genotyped in each individual fly, and **(C)** percentage of flies with genotype information for each SNP are shown. Data is split per tissue because eQTL mapping is done at the tissue level, but 593 DNA samples are used in both tissues because not only DNA, but also RNA data was available for both, head and body. Other DNA samples only have matching RNA sample derived from one tissue and are therefore only shown for that tissue.

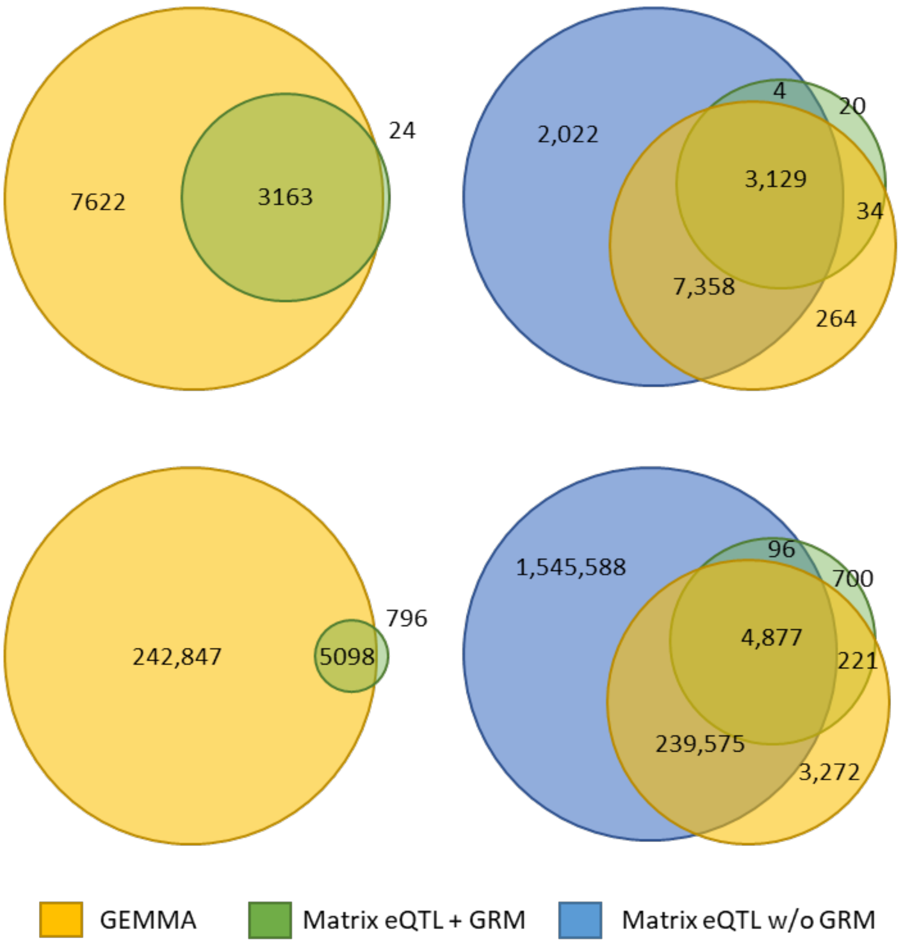


**Figure S19. Comparison of cis-eQTL (upper plots) and trans-eQTL (lower plots) results between GEMMA and Matrix eQTL**. Plots in the left show results of Matrix eQTL when the GRM is fitted as the error covariance matrix (green); 99.25% cis-eQTL and 86.5% trans-eQTL identified by Matrix eQTL are recovered by GEMMA. Plots in the right show, in addition, results of Matrix eQTL when the GRM is not included in the model, but the first five PCs of the GRM (blue). Significant eQTL are defined at 5% FDR.
